# Nucleotide diversity is a poor predictor of short-term adaptive potential

**DOI:** 10.64898/2026.01.05.697705

**Authors:** Katie L. Abson, Lillith Zijmers, Elizabeth A. Mittell, Euan A. Young, Erik Postma, Adam Eyre-Walker, Jarrod D. Hadfield

**Affiliations:** Institute of Ecology and Evolution, School of Biological Sciences, University of Edinburgh, Edinburgh, UK; School of Life Sciences, University of Sussex, Brighton, UK; Groningen Institute for Evolutionary Life Sciences, University of Groningen, Groningen, The Netherlands; Centre for Ecology and Conservation, University of Exeter, Penryn, UK

**Keywords:** Adaptive potential, Nucleotide diversity, Genetic variation, Evolvability, Conservation genetics

## Abstract

**Significance statement:** The current paradigm in conservation genetics suggests that species with the lowest molecular genetic diversity have the lowest capacity to adapt. Despite previous concern that traditional measures of molecular genetic variation are not useful predictors of adaptive potential, the conflation of genetic diversity and adaptive potential remains prevalent in both scientific literature and global policy. By combining new theory with a large dataset of genetic variation across hundreds of species, we show that molecular sequence variation is weakly related to the level of heritable variation in traits across species. This demonstrates that genetic diversity does not reliably predict adaptive potential, and highlights the urgent need to move beyond simple measures when assessing the evolutionary resilience of populations.

A capacity to adapt is essential for a population to avoid extinction in a changing world and is recognised as a global conservation priority. Adaptation requires additive (heritable) genetic variation for traits that influence survival and fecundity, but measuring this variation is difficult, particularly in species of conservation concern. Instead, molecular genetic diversity is often used to infer adaptive potential. However, previous research has cast doubt on the suitability of traditional molecular markers (allozymes and microsatellites) for this purpose given their weak relationship with heritability - a common measure of additive genetic variance. Recent advances in sequencing technology have since shifted focus towards nucleotide diversity and variation in functional regions, but their practicality for predicting adaptive potential remains debated and untested. Furthermore, heritability itself is a poor proxy for adaptive potential because it depends on environmental variance. We collated 2,113 published estimates of evolvability - a measure of additive genetic variance that avoids environmental confounding - across 193 eukaryotic species, and evaluated how well evolvability is predicted by molecular diversity. We find that microsatellite and nucleotide diversity are not significantly correlated to each other, and neither predict evolvability (nucleotide diversity explains 0.7% of interspecific differences in evolvability and doubling nucleotide diversity only corresponds to a 9.2% increase in evolvability). With new theoretical work, we show that such weak associations are expected. Together, our results suggest that simple molecular measures of genetic variation are insufficient for predicting adaptive potential and continued reliance on these metrics risks misinforming conservation management.

## Introduction

Genetic diversity has played an important role in conservation genetics since the emergence of the field more than 50 years ago, where it is often used as an indicator of a population’s genetic health (1, 2). Beyond the challenges of demographic and environmental stochasticity, small populations face an elevated risk of extinction due to the genetic consequences of their reduced size (3). Following a bottleneck, the mean fitness of a population may be reduced by inbreeding depression, where the effects of recessive deleterious variants are expressed more frequently due to increases in homozygosity. Over subsequent generations, fitness can further decline as deleterious variants stochastically increase to fixation. Finally, the genetic health of a population can be compromised by a reduction in its capacity to adapt to a change in environment. Adaptation is expected to occur mainly through changes in allele frequencies of standing genetic variation, since beneficial mutations are rare and their initial increase in frequency is slow and subject to high rates of stochastic loss (4). Therefore, it is generally believed that loss of genetic diversity limits adaptive potential and, consequently, that small populations are less robust to environmental change. Reflecting this view, 196 countries have committed to maintaining genetic diversity within and between populations to safeguard adaptive potential, a critical conservation target of the Kunming-Montreal Global Biodiversity Framework (5). However, there is no consensus on which aspects of genetic variation are essential to preserve.

Classical theory assumes that most genetic variants that will become adaptive following a change in environment currently have negligible fitness effects (6–9). Therefore, adaptive potential has traditionally been inferred from the genetic diversity of presumed neutral molecular markers, such as microsatellites or nucleotide polymorphisms in noncoding regions (2). Conversely, some argue that this approach is unhelpful and that functional variation should be prioritised instead, especially if future adaptive variants are more likely to be currently deleterious (10). In practice, identifying functionally important loci is difficult, and prioritising the diversity of a limited subset of functional loci may be counterproductive in the long term as we cannot be certain which genetic variants will be beneficial in future environments (11). Therefore, others maintain that genome-wide diversity, including neutral sites, remains crucial (11–13).

Adaptive potential is often defined loosely as ‘the capacity to adapt’, which does not provide a useful framework for assessment and conservation. A precise and practical definition of adaptive potential is the rate at which fitness can increase in one generation due to selection (14, 15), which is equal to the amount of additive (heritable) genetic variance for relative fitness, *V_A_*(*w*) (Fisher’s Fundamental Theorem of Natural Selection: 16). While *V_A_*(*w*) may not necessarily translate into observed increases in mean fitness because other factors such as density dependence can obscure or counteract genetic gains (17), it is a valuable standardized metric: populations with higher genetic variance for fitness have the capacity to evolve more rapidly in response to selection. Of course, it is impossible to know the amount of *V_A_*(*w*) that a population will have at the time of a future environmental change, and even accurately estimating current levels of *V_A_*(*w*) is difficult as it requires fitness measures for a large number of known relatives (18). For this reason, direct estimates of *V_A_*(*w*) exist for only a limited number of species (19, 20) and rarely for those of conservation concern (but see 21). Ultimately, *V_A_*(*w*) is determined by the strength of selection operating on the traits that influence survival and fecundity and their genetic (co)variances (22, 23). Therefore, if we assume that future environmental changes induce directional selection in trait space that is random and uniformly distributed, then the average additive genetic variance (*V_A_*) across traits is equal to *V_A_*(*w*) (24). This means that, *V_A_*(*w*), and therefore adaptive potential, can be approximated by the mean *V_A_*across a random sample of traits (14), though this would still require more data than is typically feasible for species of conservation concern.

Molxecular genetic diversity is an appealing proxy for adaptive potential because it is relatively easy and inexpensive to measure (2) and, under some quantitative genetic theory, genome-wide genetic diversity is expected to scale proportionally with *V_A_*(*w*) (25). However, this expectation requires strict and unrealistic assumptions of Hardy-Weinberg and linkage equilibrium, and critically, independence of allele frequency and effect (25). Under models of mutation-selection balance, allele frequencies and effects should instead be negatively related as alleles with larger deleterious effects are kept at lower frequencies (26), which is consistent with patterns observed in the genetic architecture of human complex traits (27). Therefore, in reality, the relationship between genome-wide genetic diversity and *V_A_*(*w*) could be much weaker, particularly if alleles that are currently deleterious contribute to a future adaptive response (10).

At present, the true relationship between *V_A_*(*w*) and molecular genetic diversity cannot be directly assessed because there are so few estimates of both measures within the same population or species. Instead, this relationship can be inferred by testing how strongly genetic diversity predicts *V_A_*of traits. Using this approach, two previous studies have found that the relationship between traditional molecular measures (allozyme and microsatellite diversity) and mean *V_A_* across traits is weak (14, 28). However, it is unclear whether these findings are informative to contemporary conservation genetic practices, where measuring diversity of nucleotide sites is increasingly favoured (2). Firstly, nucleotide diversity is weakly correlated with allozyme diversity across species (29). To our knowledge, no broad cross-species comparisons have been reported between nucleotide and microsatellite diversity, and the associations reported within species are inconsistent and typically based on very few populations (30–32); below, we show that there is no relationship between microsatellite and nucleotide diversity across a larger sample of species (Fig. 3). Secondly, both of these earlier studies (14, 28) assessed how well genetic diversity predicts variance-standardised *V_A_*, *heritability* or *h*^2^. Although it is the most widely reported standardisation of *V_A_*, *h*^2^ is a poor comparative measure due to its dependence on environmental and non-additive genetic variance (33, 34), which may contribute to the weak correlation between heritability and molecular diversity (28) or population size (35). Instead, the mean-standardised *V_A_*, *evolvability* or *I_A_*, has been proposed as a more suitable quantitative genetic measure of adaptive potential (33, 34). *I_A_* quantifies the maximum proportional change in the trait mean per generation under a standardised strength of selection (equivalent to selection acting on fitness itself), and unlike *h*^2^, *I_A_* is unaffected by non-additive genetic and environmental variance. Like the different measures of molecular diversity, *I_A_* and *h*^2^ are only weakly correlated with each other, particularly in natural populations (33, 34, 36, 37). Crucially, this means that the weak relationships that have previously been reported between allozyme or microsatellite diversity and heritability do not necessarily imply that the nucleotide diversity is a poor measure of adaptive potential.

Limited data availability previously prevented the assessment of whether nucleotide diversity predicts a species’ mean evolvability of traits (14) and the nature of the relationship between molecular genetic variation and evolvability remains a key question in evolutionary biology (38). In light of the global recognition of the importance of adaptive potential, and ongoing debate regarding the utility of different genetic measures for predicting it, addressing this question is now essential. Here, we compile thousands of published estimates of quantitative (evolvability, *I_A_* and heritability, *h*^2^) and molecular (pairwise nucleotide diversity, *π* and microsatellite expected heterozygosity, *H_e_*) genetic variation across 246 eukaryotic species (Table 1). We evaluate how reliably molecular genetic measures predict adaptive potential, first by quantifying the observed relationships between molecular and quantitative genetic variation across species, and then using theoretical models to assess the expected strength of these associations under various evolutionary conditions. We additionally test whether quantitative genetic variation differs among Red List categories, in line with recent discussions on the potential inclusion of genetic data for evaluation of species’ IUCN Red List status (39).

**Table 1:**
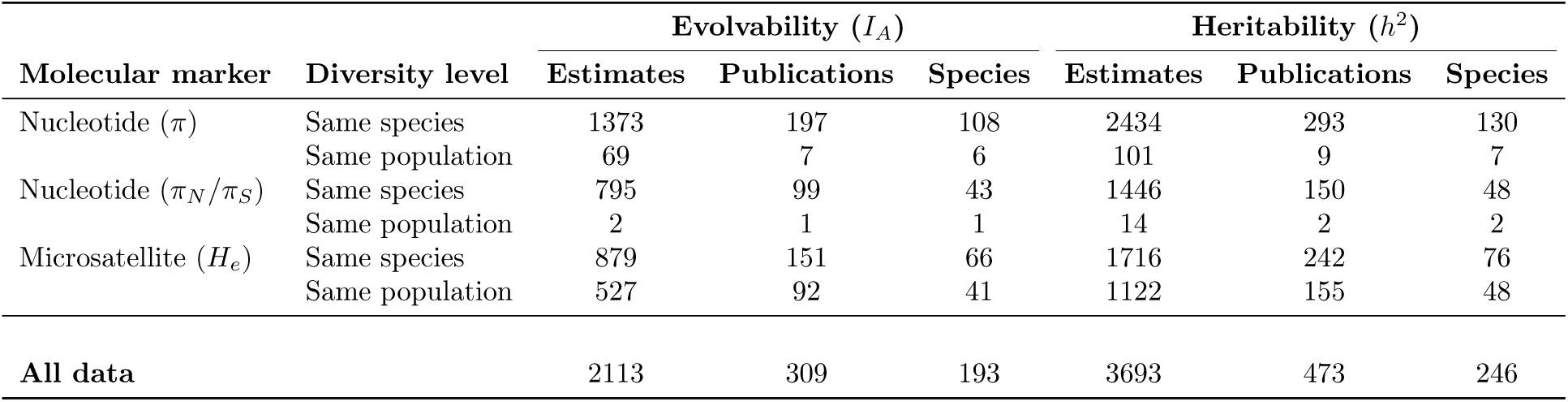
Summary of sample sizes for main analyses of mean-scaled additive genetic variance (evolvability) and variance-scaled additive genetic variance (heritability). Sample sizes for the univariate analyses were dependent on the availability of molecular diversity estimates (*π* = pairwise nucleotide diversity, *π_N_ /π_S_* = ratio of nonsynonymous to synonymous pairwise nucleotide diversity, *H_e_* = expected heterozygosity) from the same species; the number of estimates where a corresponding molecular measure was available from the same population is indicated. Bivariate analyses used all available quantitative genetic data.

| Molecular marker | Diversity level | Evolvability ($I_A$) | | | Heritability ($h^2$) | | |
| --- | --- | --- | --- | --- | --- | --- | --- |
|  |  | Estimates | Publications | Species | Estimates | Publications | Species |
| Nucleotide ($\pi$) | Same species | 1373 | 197 | 108 | 2434 | 293 | 130 |
|  | Same population | 69 | 7 | 6 | 101 | 9 | 7 |
| Nucleotide ($\pi_N/\pi_S$) | Same species | 795 | 99 | 43 | 1446 | 150 | 48 |
|  | Same population | 2 | 1 | 1 | 14 | 2 | 2 |
| Microsatellite ($H_e$) | Same species | 879 | 151 | 66 | 1716 | 242 | 76 |
|  | Same population | 527 | 92 | 41 | 1122 | 155 | 48 |
| **All data** |  | 2113 | 309 | 193 | 3693 | 473 | 246 |

## Results

### Nucleotide diversity vs. evolvability

To investigate the relationship between genome-wide nucleotide diversity, *π̂*, and *true* average trait evolvability, *I_A_*, we compiled 1,373 published estimates of evolvability from 108 species for which measures of putatively neutral nucleotide diversity were also available. Using univariate linear mixed models that control for phylogenetic relatedness and differences in estimation between evolvability estimates, we found no meaningful association between *ln*(*I_A_*) and *ln*(*π̂*) (Fig. 1; Fig. 2). Estimated parameters are summarised by their posterior median and 95% credible intervals. The slope of the regression was *β* = 0.127 [−0.202 – 0.410] (P=0.397) and despite substantial variation in *ln*(*I_A_*) between-species (standard deviation of 1.533 [0.649 – 2.426]), only *R*^2^ = 0.7% [−3.0 – 9.5] of this could be predicted by *ln*(*π̂*) (note we carry the sign in the *R*^2^ value). We also calculated 2*^β^*, a measure of the relationship that is independent to the amount of interspecific variaton in *ln*(*I_A_*) (see methods), which suggests that doubling *π̂* would only increase mean evolvability by 9.2% [−13.1 – 32.8]. There was substantial variation *ln*(*Î_A_*) within-species, some of which can be explained by methodological or trait differences (details in SI section 2). An equivalent bivariate model, which leverages information from 85 additional species with available estimates of *I_A_* but not *π̂*, estimated a similarly weak association between *ln*(*I_A_*) and *ln*(*π̂*) but with wider credible intervals (*β* = −0.043 [−0.606 – 0.489], P=0.872, 2*^β^* = −2.9% [−36.9 – 37.7] and *R*^2^ = −0.2% [−20.3 – 15.8]). *ln*(*π̂*) was also weakly predictive of species’ average heritability, *h*^2^ (SI; Fig. S3).

**Figure 1:**
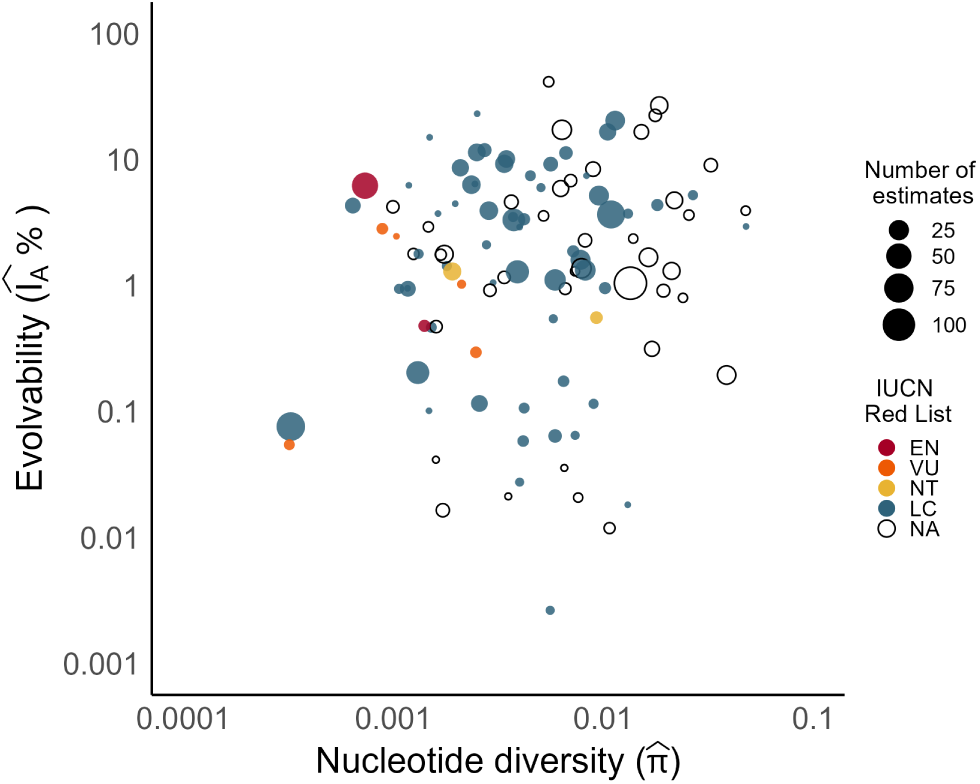
Average *measured* mean-scaled additive genetic variance (evolvability, *Î_A_*) versus neutral pairwise nucleotide diversity (*π̂*) assayed in the same species (n=108), excluding non-positive estimates. Both measures are plotted on log_10_ scale, though data were log*_e_*-transformed for analyses. The size of the points shows the number of estimates over which the average evolvability is calculated and the colour shows the IUCN Red List category for that species, where EN = Endangered, VU = Vulnerable, NT = Near threatened, LC = Least concern, and NA = Not classified.

**Figure 2:**
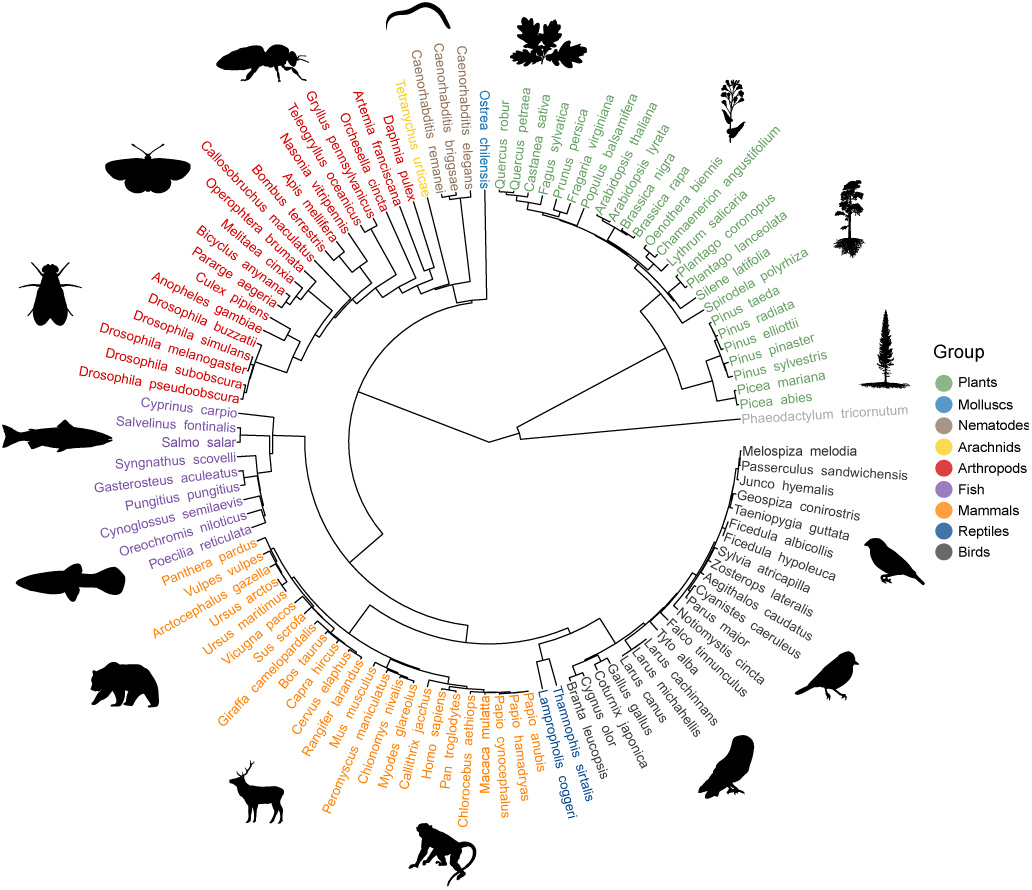
Phylogeny used in the univariate analysis of evolvability (*I_A_*) with nucleotide diversity (*π̂*) as a predictor, visualised using ggtree v3.10.1 (40) and rphylopic v1.5.0 (41).

Some authors suggest that functional variation is more critical for conservation of adaptive potential (10). Nonsynonymous diversity, *π_N_*, is a relatively easy measure of putatively functional (selected) variation but is strongly correlated with neutral *π* (42). We therefore repeated the univariate and bivariate analyses, substituting genome-wide diversity *ln*(*π̂*) for the ratio of nonsynonymous to synonymous diversity, *ln*(*π̂_N_ /π̂_S_*), as an alternative simple metric which depends on both effective population size, *N_e_*, and the average strength of selection (43). The credible intervals were wider than those of the equivalent analysis of neutral nucleotide diversity, but point estimates were similarly weak. The slope of the regression was *β* = 0.086 [−0.599 – 0.753] (P=0.802), therefore doubling *π̂*_N_ /π̂*_S_* only corresponds to an increase in evolvability by 6.1% [−37.5 – 64.6], and 0.1% [−7.0 – 23.0] of variation in *ln*(*I_A_*) could be predicted by *ln*(*π̂_N_* /*π̂_S_*). The equivalent bivariate model estimated a stronger association between *ln*(*I_A_*) and *ln*(*π̂_N_ /π̂_S_*) but with wider credible intervals (*β* = 0.621 [−0.297 – 1.386], P=0.160, 2*^β^* = 53.8% [−18.6 – 161.4] and *R*^2^ = 11.8% [−1.0 – 37.0]). *ln*(*π̂_N_ /π̂_S_*) was also weakly predictive of species-mean *h*^2^ (SI; Fig. S3).

### Microsatellite diversity vs. evolvability

Although used less frequently, microsatellites are still popular markers of choice in conservation genetics (2). We estimated the correlation between mean nucleotide *π* and mean microsatellite expected heterozygosity, *H_e_*, across 57 species and found it to be weak and non-significant (*r* = −0.031 [−0.292 – 0.226], P=0.836; Fig. 3). Therefore, microsatellites and nucleotides may differ in their ability to predict adaptive potential.

**Figure 3:**
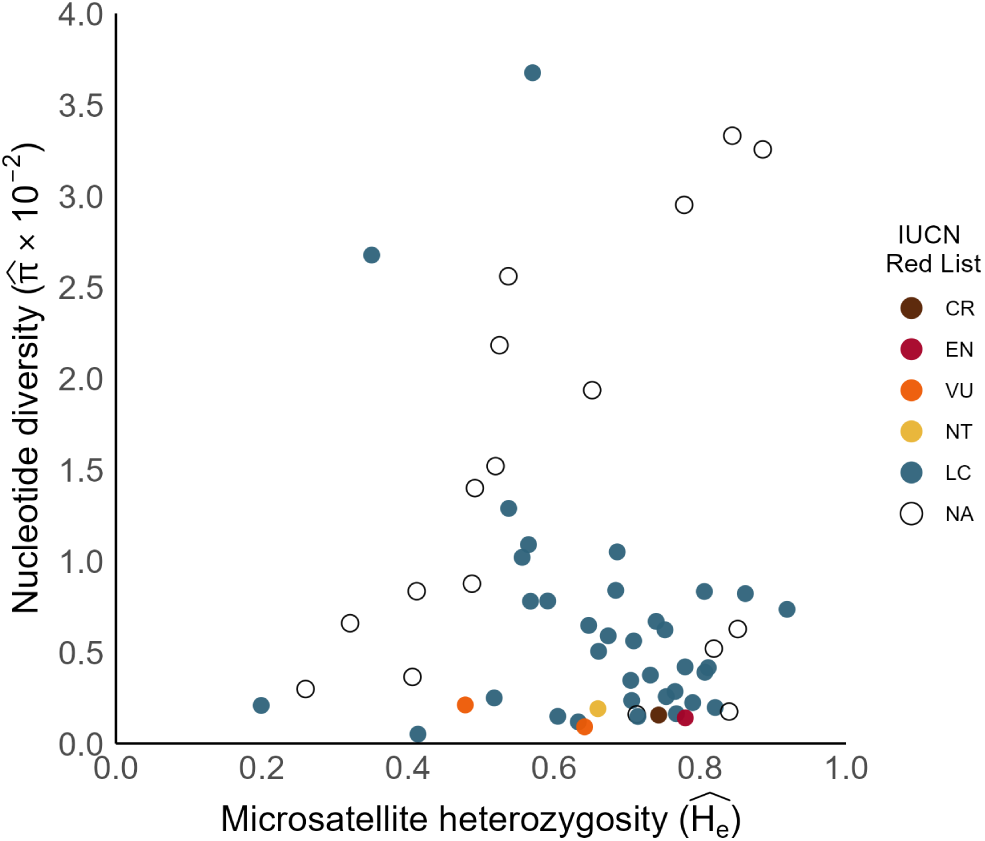
Mean neutral pairwise nucleotide diversity (*π̂*) versus microsatellite expected heterozygosity (*Ĥ_e_*) across 57 eukaryotic species. Colour shows the IUCN Red List category for the species, where CR = Critically endangered, EN = Endangered, VU = Vulnerable, NT = Near threatened, LC = Least concern, and NA = Not classified.

Previous work has shown that microsatellite diversity is a weak predictor of *h*^2^ (14). However, if *I_A_* is a better proxy for adaptive potential and *h*^2^ and *I_A_* are only weakly correlated, microsatellite diversity may be a better predictor of adaptive potential than this earlier study suggests. A weak relationship between *h*^2^ and *I_A_* has previously been shown across traits (33, 34, 36) - a result we also observe here with a bivariate model of 1,976 *h*^2^ and *I_A_*estimates (*r* = 0.135 [0.077 – 0.196], P*<*0.5*×*10*^−^*^3^; Fig. S2). We additionally show that across 183 species, average levels of *h*^2^ and *I_A_*are also weakly and not significantly correlated (*r* = 0.130 [−0.085 – 0.366], P=0.271). Nevertheless, when we used the univariate model described above with *ln*(*π̂*) substituted for *ln*(*Ĥ_e_*), the regression of *ln*(*I_A_*) on mean *ln*(*Ĥ_e_*) was also found to be weak (*β* =-0.535 [−1.308 – 0.344], P=0.237), therefore doubling *Ĥ_e_* may reduce mean evolvability by −31.0% [−66.2 – 15.8]), but credible intervals are wide. The *R*^2^ was effectively zero (−0.9% [−8.7 – 1.3]).

Since our dataset contains more data for species with both *h*^2^ and microsatellite diversity than was previously available (14) - specifically, 48% of *h*^2^ estimates were not included in the previous study and a further 13 species are represented - we also quantified the regression of *h*^2^ on *ln*(*Ĥ_e_*). Consistent with the findings of (14), initial models found that *ln*(*Ĥ_e_*) was predictive of *h*^2^ (*β* =0.088 [0.065 – 0.109], P*<*0.5*×*10*^−^*^3^ and *R*^2^ = 3.9% [0.3 – 8.8]), but this association was largely driven by a single study on *Arabidopsis thaliana* (44). When data from this study were excluded, the estimated relationship was weaker with wider credible intervals (*β* =-0.033 [−0.130 – 0.062], P=0.512 and *R*^2^ = −0.5% [−10.4 – 4.4]). Details of these additional analyses are provided in the supplementary materials and sample sizes are shown in Table 1.

### Theoretical expectations

For neutral traits at equilibrium a one-to-one relationship between *ln*(*V_A_*) and *ln*(*π*) is expected. Under non-equilibrium conditions, it has been suggested that the relationship may be less than one-to-one due to the different rates of mutational input for quantitive traits and nucleotide sites (45, 46). However, we show that even under non-equilibrium conditions a one-to-one relationship is expected under neutrality if *V_A_* and *π* have been at equilibrium at some point previously (Fig. S6). In contrast, for traits under stabilising selection, the relationship at equilibrium is expected to be weak unless *N_e_* is very small and/or selection on the underlying loci is very weak (Figs. S4 and S5). Under non-equilibrium conditions, stabilising selection on a trait can speed up the rate at which *V_A_* equilibrates, further decoupling *V_A_* from *π*, although this effect is expected to be relatively weak for polygenic traits (Fig. S7). See SI for full details.

### Quantitative genetic variation vs. IUCN Red List

Finally, we tested whether quantitative genetic variation differed among Red List categories by including the species’ IUCN Red List status as a predictor (but omitting *π̂*). We did not find significant differences in mean *I_A_* or *h*^2^ across IUCN Red List categories (*I_A_*: *χ*^2^*_4_* = 0.264, P=0.992, Table S9; *h*^2^: *χ*^2^*_4_* = 6.279, P=0.179, Table S10). Because estimates for threatened species were limited (Table S11, Fig. 1), we repeated the analysis with higher risk categories grouped. Mean *I_A_* was 3.5% [−54.8 – 84.8] (P=0.911) higher and mean *h*^2^ was 0.08 [−0.01 – 0.16] (P=0.053) higher in species of conservation concern. However, given the credible intervals on these differences remain wide, we consider this an open question.

## Discussion

Previous studies have shown that both microsatellite and allozyme diversity are poor predictors of heritability (14, 28). However, these findings may give little insight into whether conserving nucleotide diversity can maintain adaptive potential, as mandated by the UN’s landmark biodiversity agreement (5), because these traditional markers are weakly correlated with nucleotide diversity (Figs. 3 and 4) and heritability itself may be an unreliable proxy. To assess whether contemporary molecular measures of genetic variation can predict adaptive potential, we estimated the strength of the relationship between nucleotide diversity and evolvability (*I_A_*, mean-standardised *V_A_*) across 108 species. Our results suggest that nucleotide diversity is a poor predictor of evolvability across species, with the best estimates showing only 0.7% of variation in *ln*(*I_A_*) can be explained by variation in nucleotide diversity and that *I_A_* would only increase by 9.2% if nucleotide diversity were doubled. Similarly weak relationships were found across the different measures of molecular and quantitative genetic variation (Fig. 4), suggesting that simple assays of molecular genetic diversity are unlikely to be useful for inferring a species’ adaptive potential.

**Figure 4:**
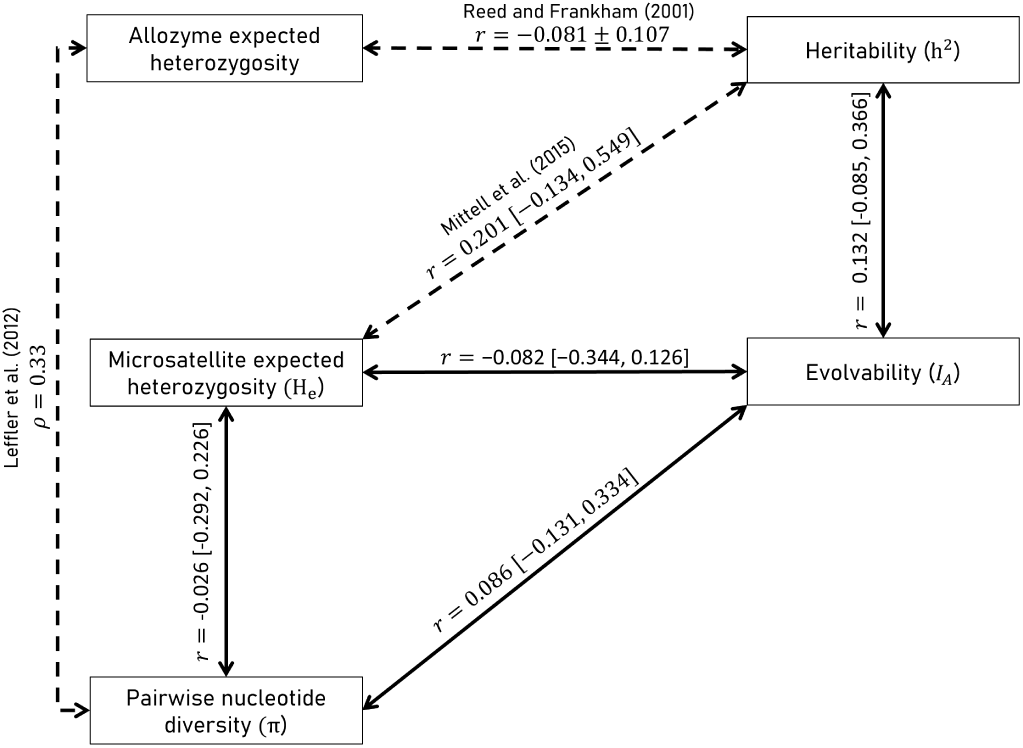
Correlations between measures of molecular (left: allozyme expected heterozygosity; microsatellite expected heterozygosity, *H_e_*; and pairwise nucleotide diversity, *π*) and quantitative (right: average heritability, *h*^2^; and average evolvability, *I_A_*) genetic variation from analyses using data from over 20 species. Solid lines show estimates from this study and dotted lines show results reported in previous studies (14, 28, 29). Values show the Spearman rank (*ρ*) or Pearson (*r*) correlation coefficient ± the standard error or 95% credible intervals in brackets. The correlation coefficient is not provided explicitly in 14, so we calculated it directly from their published population-level model which excluded estimates from 44. Note that 28 report the correlation between allozyme diversity and *measured* average heritability, *ĥ^2^*. See 33 for the estimated correlation between heritability and evolvability of individual traits.

A strong one-to-one relationship between *ln*(*V_A_*) and *ln*(*π*) is only expected if quantitative traits are neutral. Our estimate of the slope is considerably less than one (*β* = 0.127 [−0.202 – 0.410]) which is instead consistent with quantitative traits being subject to stabilising selection and weakly deleterious variants making a substantial contribution to existing *V_A_*. Although classic conservation genetic theory assumes that effectively neutral genetic variation will underpin future adaptive responses, forming the basis of rules such as the *N_e_*=500 guideline (8), under a trait-based model of adaptation, it is these currently deleterious variants that are likely to become advantageous when the optimal phenotype changes (10, 16, 47). The degree to which stabilising selection can decouple the relationship between *ln*(*V_A_*) and *ln*(*π*) depends critically on *N_e_*. At equilibrium in high *N_e_* populations, the relationship between *V_A_*and *π* is driven by their shared dependence on the mutation rate, and so the relationship is likely weak if most of the variation in *π* is driven by differences in *N_e_*. Conversely, if *N_e_* is very small and/or selection at quantitative trait loci is very weak, the relationship converges to the neutral expectation of a one-to-one relationship. Whilst species of conservation concern might be expected to have lower *N_e_* and therefore a stronger *ln*(*V_A_*)-*ln*(*π*) relationship than the species surveyed here, our dataset spans a range of nucleotide diversities typical of such species (e.g., Fig. 1 in 10). The absence of any detectable relationship within this range indicates that the weak association between *π* and *I_A_* is not simply an artifact of sampling many non-threatened taxa (11). Additionally, in using the mean *I_A_*of traits as a proxy for *V_A_*(*w*), we implicitly assume that traits are equally likely targets of future selection. It is probably more realistic that most future adaptive traits are those already closely associated with fitness and therefore under stronger selection than the average trait sampled here. If anything, this would further weaken the dependence of *V_A_*(*w*) on *π*. Finally, although few species had available estimates of both evolvability and nucleotide diversity from the same population (Table 1), using diversity estimates assayed in different populations of the same species should not strongly affect estimated genome-wide diversity since there is relatively low variation in nucleotide diversity among populations (14). Therefore, the relationship between *I_A_* and *π* is unlikely to be usefully stronger at the population-level.

While a strong relationship between *ln*(*V_A_*) and *ln*(*π*) might be expected in very small populations at drift-selection-mutation equilibrium, it has been repeatedly suggested that departures from equilibrium may explain any decoupling between these measures (14, 45, 46, 48). Although genetic drift reduces both *V_A_* and *π* by a fraction of 1*/*2*N_e_*each generation (49), mutation replenishes quantitative genetic variation far faster (10*^−^*^3^ - 10*^−^*^2^ per trait per generation, 50) than nucleotides (10*^−^*^9^ - 10*^−^*^8^ per site per generation, 51). Consequently. it has been suggested that moderately recent bottlenecks or expansions may render *π* as an out-of-date snapshot of a population’s adaptive potential (14, 45, 46, 48). We show that this idea is only partly true: under neutrality a population only has to come to equilibrium once, and in the generations thereafter the relationship between *ln*(*V_A_*) and *ln*(*π*) will be close to one-to-one, irrespective of whether the population remains in equilibrium or experiences a change in population size. External perturbations to allele frequencies (e.g. a translocation event) are required to cause the relationship to deviate from one-to-one. Conversely to previous suggestions (14, 48), this implies that the higher mutation rate of microsatellites may not offer any appreciable advantage over nucleotides in bottlenecked populations. Consistent with this, we find that microsatellite diversity is also weakly predictive of quantitative genetic variation, although this may be compounded by ascertainment bias for highly polymorphic microsatellite markers (14). In non-neutral models, however, *V_A_* equilibrates faster as the strength of selection on the underlying loci increases (52), which can cause a decoupling from the genetic diversity of loci under weaker selection. Nevertheless, this effect diminishes as the trait becomes more polygenic because selection acting on individual loci then becomes small (52). So for many traits, including fitness, the lack of relationship between *ln*(*V_A_*) and *ln*(*π*) is unlikely to be solely driven by departures from equilibrium, though it is unclear how robust these expectations are to more realistic patterns of dominance, epistasis, linkage and pleiotropy. With this in mind, we tested whether a metric which integrates information for putative selected diversity, *π_N_ /π_S_*, would be a better predictor of *V_A_* than *π* since the loci contributing to *π_N_* should have selection coefficients more comparable to quantitative trait loci. However, the relationship between *π̂_N_ /π̂_S_* and measures of *V_A_* could not be estimated with sufficient precision, and while *π_N_ /π_S_* is expected to depend on the mean strength of selection (and *N_e_*) (43), there may be little variation in the mean strength of selection across species, though this has only been characterised in a few taxa (e.g. 53).

While the weak association between nucleotide diversity and *V_A_* does not necessarily mean that genetic diversity is a poor predictor of overall genetic health, the evidence that genetic diversity predicts other genetic components of extinction risk also remains mixed (10, 11) and techniques have moved towards more advanced predictions based on genomic annotation and conservation (54, 55). Given that there is substantial inter-specific variation in trait evolvability (see also 56), consistent with differences in *V_A_*(*w*) between populations and/or species (19), predicting this variation remains a priority and it is an open question whether newer functionally-informed methods could also improve prediction of adaptive potential. However, if weakly deleterious variants contribute to future adaptive responses, a possible negative relationship between segregating load and adaptive potential could complicate assessment of conservation priority.

By compiling a large dataset of genetic variation across a diverse range of taxa, and explicitly accounting for methodological variation among published estimates of evolvability, we show that nucleotide diversity fails to predict a species’ *true*level of quantitative genetic variation – the necessary component for a response to selection. This echoes earlier work that found weak relationships between other measures of molecular diversity and quantitative genetic variation (14, 28). Given our definition and proxy for adaptive potential (the capacity for an immediate response to directional selection following a change in environment), these results highlight that existing molecular genetic proxies are unsuitable for predicting the capacity of a species to adapt to changes in their environment. Longer-term responses will depend on both *N_e_* during the period of directional selection (57) and new mutations (58). The degree to which *π* predicts this longer-term response depends on the accuracy with which *π* predicts future *N_e_*, which may be poor in rapidly declining populations (59). Overall, the recent global emphasis on conserving adaptive potential is encouraging, but the absence of a widely accepted formal definition and reliance on weak proxies impedes effective population assessment and management.

## Materials and methods

### Data

2,040 new estimates of heritability (*h*^2^) and 1,145 new estimates of evolvability (*I_A_*, or the coefficient of additive genetic variance, 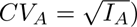 were combined with 1,653 *h*^2^ and 987 *I_A_* or *CV_A_* estimates from two previous meta-analyses (14, 60), producing a final dataset that included *h*^2^ estimates from 473 publications representing 246 species, and *I_A_* estimates from 309 publications representing 193 species. New estimates were obtained by searching Web of Science for studies published between 1992 and 2022 in the journals ‘Journal of Evolutionary Biology’, ‘Evolution’, ‘Heredity’, and ‘Proceedings of the Royal Society B’ with the topics ‘heritability’ and ‘evolvability’. If evolvability was not reported, we calculated it from its component parts where possible. For each estimate, we recorded trait type (condensed classification of 33: morphological, physiological, behavioural, life history, fitness), method of estimation (clonal, full/half-sib model, mid/single-parent-offspring regression, animal model, realised selection response), dimensional classification (linear, quadratic, cubic, meristic, time, other) and model structure (the number of fixed/random effects). Because evolvability estimates are not suitable for all traits (33, 61) and often contain errors (62), we checked all evolvability estimates for inclusion, then kept only one estimate per-trait per-population (usually estimates from the largest sample) to avoid non-independence.

Single- and multi-locus estimates of autosomal gene diversity compiled by 14 provided the basis for our molecular dataset. This included 2,788 microsatellite expected heterozygosity (*H_e_*) estimates in 71 species and 1,777 pairwise nucleotide diversity (*π*) estimates at putatively neutral sites (silent, synonymous, 4-fold degenerate, noncoding and intronic sites) in 45 species. To increase overlap between the molecular and quantitative genetic datasets, further *π* estimates were sourced from other meta-analyses (42, 51, 63–65) and by searching Web of Science with the keywords ‘genetic diversity’ or ‘nucleotide diversity’ and the Latin species name. We additionally collected genome-wide diversity estimates for 47 species from the Tree of life data portal (66). Although these estimates are based on a single genome per species, such values typically provide reliable approximations of nucleotide diversity calculated from larger samples (Fig. S1). To obtain estimates of diversity at ‘functional’ (presumed selected) sites, we revisited the original publications and recorded estimates of nonsynonymous or 0-fold degenerate pairwise nucleotide diversity (*π_N_*). The final dataset included 4,202 single- and multi-locus microsatellite *H_e_*estimates (104 species), 3,170 *π* estimates (192 species), and 764 *π_N_* estimates (87 species). The relevant data structures for the analyses of quantitative and molecular genetic variation are summarised in Table 1.

### Analyses

All statistical analyses involved Bayesian linear mixed models fitted using the package MCMCglmm v2.35 (67) in R v4.4.2 (68). Scaled (by 1,000) *F*_1,1_ priors were used for all random-effect variance components and an inverse gamma prior, with shape and scale equal to 0.002, was used for the residual variance component. Normal priors with zero mean and large variances (10^8^) were used for the fixed effects. The MCMC chains were run for 500,000 iterations with a burn-in period of 100,000, sampling every 200 iterations. Significance was assessed using pMCMC values (69) or a Wald test for omnibus tests of multi-category factors where the posterior means and covariance matrix of the effects were used. A threshold of 0.005 was used to determine significance and values between 0.005-0.05 were considered suggestive (70).

### Nucleotide diversity vs. evolvability

As theory predicts a proportional relationship between *V_A_* and genetic diversity (see SI section 3), both evolvability and nucleotide diversity were log transformed prior to analysis. *ln*(*π̂*) was fitted as a covariate with regression coefficient *β*. We do not adjust for methodological heterogeneity among *π̂* estimates as this variation reflects the practical challenge of inferring *V_A_* from empirical data. True variation in *ln*(*I_A_*) among species, after accounting for *ln*(*π̂*), was estimated by fitting *phylogenetic* and *non-phylogenetic* species effects as random. The phylogenetic covariance structure was assumed to be proportional to the amount of time two species have shared ancestry, based on a phylogeny constructed from published divergence-time estimates (Fig. 2; 71). The tree was scaled to unit length such that the estimated variance in the phylogenetic (*V*_*ln*(*IA*):Phy_) and non-phylogenetic (*V*_*ln*(*IA*):S_) species effects sum to give the total remaining between-species variation in *ln*(*I_A_*). Differences in *ln*(*I_A_*) estimation were accounted for with fixed effects for *type of relative* and the number of *fixed* and *random effect terms*. Trait differences were modelled with fixed effects for *trait category* and *trait dimension* and a random effect for *trait* identity. A random *publication* effect was fitted to capture remaining differences among studies not explained by these predictors.

From the model, two measures of the relationship between *π̂* and *I_A_* were calculated. First, the coefficient of determination,

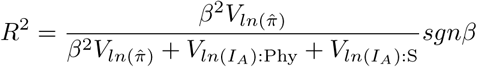

where the denominator is the total between-species variance in *ln*(*I_A_*) and *V*_*ln*(π̂)_ is the variance in log-transformed nucleotide diversity estimates such that the product *β*^2^*V*_*ln*(π̂)_ is the variation in *ln*(*I_A_*) owing to variation in *ln*(*π̂*). The sign of the regression coefficient is carried to allow for negative *R*^2^ values (as in 14). As with *h*^2^, *R*^2^ is prone to a ‘rubber ruler’ effect (61) as it is dependent on the amount of variation in *I_A_*. This means that a high *R*^2^ would only indicate that molecular measures are useful if there is sufficient variation in *I_A_* to expect meaningful differences in adaptive potential between species. Therefore, we also report 2*^β^*, the expected proportional change in mean *I_A_* if *π̂* is doubled, which is independent of interspecific variation in *I_A_*.

In addition, we fitted an equivalent bivariate model with both *ln*(*Î_A_*) and *ln*(*π̂*) as response variables, allowing us to make use of all available quantitative genetic data, including species for which *π̂* was missing but available for a close relative. Details are provided in the supplementary materials.

Nonynonymous diversity (*π_N_*) is strongly correlated with neutral (synonymous) genetic diversity (42) meaning that *π_N_* alone has limited scope to better predict evolvability (14). Instead, we used the ratio of nonsynonymous (or 0-fold degenerate) and synonymous (or 4-fold degenerate) diversity (*π_N_ /π_S_*) as a measure of functional variation. The relationship between *ln*(*I_A_*) and *ln*(*π̂_N_ /π̂_S_*) was assessed using the same univariate and bivariate model structures described above, with *ln*(*π̂_N_ /π̂_S_*) replacing *ln*(*π̂*). All other predictors and MCMC specifications remain unchanged.

### Microsatellite diversity vs. evolvability

Before assessing the relationship between microsatellite diversity and evolvability, we first compared traditional and contemporary measures of genetic variation using two bivariate models. For molecular genetic variation, the responses were species-average *π* and species-average microsatellite *H_e_*, fitted across 57 species with available estimates of both measures. For the quantitative genetic measures, the responses were 1,976 estimates of *h*^2^ and *I_A_*in 183 species across the traits for which both quantitative genetic measures were available and appropriate. In the quantitative genetic model, *Species* was included as a random effect, allowing the (co)variance between *h*^2^ and *I_A_* to be partitioned into among-species and among-traits (within-species) components. For both models, the correlation coefficient (*r*) was calculated from posterior samples of the (co)variance matrices; for the quantitative genetic model, correlations were estimated separately at the species- and trait-level.

We then assessed the relationship between *ln*(*I_A_*) and microsatellite *ln*(*Ĥ_e_*) using the same univariate and bivariate model structures described for the analyses of nucleotide diversity, with *ln*(*Ĥ_e_*) replacing *ln*(*π̂*) as a predictor. All other predictors and MCMC specifications remain unchanged.

### Theoretical expectations

To support the empirical analyses of molecular versus quantitative genetic variation, we theoretically explored the expected relationship between *V_A_* and genetic diversity under a range of models for drift-selection-mutation, both at equilibrium and out of equilibrium. To achieve this we derived the partial derivatives of *ln*(*V_A_*) with respect to the logged mutation rate, *µ*, and *N_e_*. From these we approximated the expected regression of *ln*(*V_A_*) on *ln*(*π*) *≈ ln*(4) + *ln*(*µ*) + *ln*(*N_e_*) as

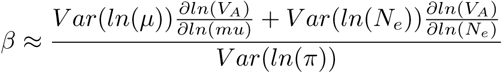

under the assumption that *ln*(*µ*) and *ln*(*N_e_*) are not strongly correlated. If they are negatively correlated (72) the regression will be shallower than this approximation predicts. See SI for details.

### Quantitative genetic variation vs. IUCN Red List

Species’ Red List status was recorded from the IUCN website (73). Univariate models of *ln*(*I_A_*) and *h*^2^ were refitted with Red List category (least concern, near threatened, vulnerable, endangered or critically endangered) as a predictor, excluding molecular genetic diversity. Given the paucity of estimates for species of conservation concern according to the Red List, both models were repeated with a condensed classification of extinction risk in which all at-risk categories were grouped into a single ‘conservation concern’ group and compared to species of ‘least concern’. The sample size per category is given in Table S11.

## Acknowledgements

We thank Lucie Bergeron for sharing nucleotide diversity data and Thomas Hansen, David Houle and Christophe Pélabon for insightful discussion regarding evolvability estimates. K.L.A was supported by funding from NERC through an E4 DTP studentship (NE/S007407/1).

## Author contributions

E.A.M, E.A.Y and E.P provided data. K.L.A, L.Z and E.A.M collected new data. K.L.A and L.Z compiled and curated data. K.L.A, L.Z, A.E.W and J.D.H developed the statistical analyses and J.D.H developed the theoretical analyses. K.L.A and J.D.H wrote the initial draft and all authors contributed to the editing of the manuscript.

