## Supplementary Information for "Nucleotide diversity is a poor predictor of short-term adaptive potential"

January 5, 2026

### 1 Data

#### Quantitative genetic variation

Fewer estimates of evolvability were available compared with heritability since it is less frequently reported and because evolvability is not suitable for all traits (1, 2). In many cases where evolvability had not been reported, we were able to calculate it using the formula $I_A = V_A/\bar{X}^2$, where $V_A$ is the additive genetic variance and $\bar{X}$ is the trait mean (note that only the denominator is squared). If neither $V_A$ or the trait mean were reported, an evolvability estimate was still able to be calculated if an estimate of both the heritability, $h^2$ and the coefficient of variation $CV = \sqrt{V_P}/\bar{X}$, where $V_P$ is the phenotypic variance, were available.

For each estimate, we recorded additional information relating to the choice of trait and methodology. We recorded the trait type using a condensed classification of 2: morphological, physiological, behavioural, life history and fitness (lifetime breeding success), since both heritability and evolvability have been shown to vary among these categories (2–5). Since evolvabilities depend on the dimension of a trait (2), we further classified estimates as: linear, quadratic, cubic, meristic, time and other. To deal with methodological differences, we recorded the method of estimation: clonal, full/half-sib model, mid/single-parent-offspring regression, animal model and realised selection response, since estimation based on different types of relatives or realised changes in the trait vary in the extent to which $V_A$ is biased (6). Finally we recorded the information relating to model structure (the number of fixed/random effects), since this can affect how much trait variation is estimated and partitioned among additive versus non-additive components (7).

##### Nucleotide diversity

In addition to the estimates compiled by previous meta-analyses of putatively neutral genetic diversity (4, 8–12), we obtained whole-genome diversity ($\hat{\pi}_{ToL}$) estimates for 47 species from the Tree of life (ToL) data portal (13). Since these estimates are derived from a k-mer distribution (14) of a single genome per species, we first assessed how they compared to published pairwise nucleotide diversity estimates ($\hat{\pi}$) across the 15 species for which we had both measures. A simple linear regression of $\hat{\pi}_{ToL}$ on $\hat{\pi}$ indicated a strong relationship between the two measures (estimate followed by 95% confidence intervals: intercept = 0.005[-0.116×$10^{-5}$ – 0.010]; slope = 0.657[0.377 – 0.936]; $R^2 = 0.664$) (Figure 1). If both $\hat{\pi}_{ToL}$ and $\hat{\pi}$ are estimates of the same underlying quantity, $\pi$, then deviations of $\hat{\pi}$ from $\pi$ will result in an attenuated regression slope of $\widetilde{\beta} = R^2_{\pi,\hat{\pi}}\beta$ where $\beta$ is the regression had $\pi$ been used as a predictor. Earlier work (4) estimated $R^2_{\pi,\hat{\pi}} = 0.79$ such that our estimate of the slope $\widetilde{\beta}$ is consistent with both $\hat{\pi}_{ToL}$ and $\hat{\pi}$ being estimates of the same underlying quantity and the true regression, $\beta$, being equal to one (i.e our estimated slope, $\widetilde{\beta}$, does not differ significantly from 0.79). Therefore, we included the ToL diversity estimates in our final nucleotide diversity ($\hat{\pi}$) dataset.

#### 2 Empirical analyses

##### Univariate models

As outlined in the main text, all statistical analyses involved Bayesian linear mixed models fitted using the package MCMCglmm v2.35 (15) in R v4.4.2 (16). Scaled (by 1,000) $F_{1,1}$ priors were used for all random-effect variance components and an inverse gamma prior, with shape and scale equal to 0.002, was used for the residual variance component. Normal priors with zero mean and large variances ($10^8$) were used for the fixed effects. The MCMC chains were run for 500,000 iterations with a burn-in period of 100,000, sampling every 200 iterations. Significance was assessed using pMCMC values (17) or a Wald test for omnibus tests of multi-category factors where the posterior means and covariance matrix of the effects were used. Posterior

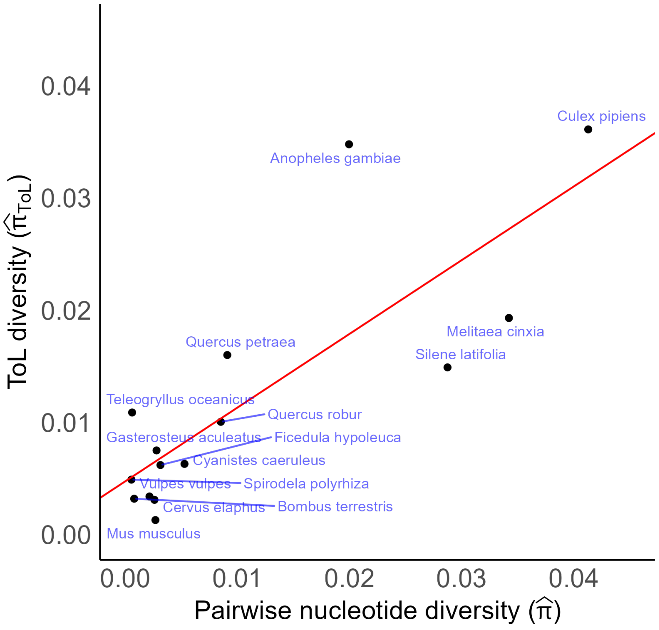


Figure 1: Estimates of diversity ($\hat{\pi}_{ToL}$) from the Tree of Life (ToL) data portal against published estimates of pairwise nucleotide diversity ($\hat{\pi}$) based on multiple individuals, across 15 species. The regression line (slope = 0.657) is shown in red.

distributions are summarised by their median (18) and 95% credible intervals. A threshold of 0.005 was used to determine significance and values between 0.005-0.05 were considered suggestive (19).

##### Molecular genetic diversity vs. Evolvability

The relationship between a population's additive genetic variance, $V_A$, and genetic diversity is expected to be proportional under some evolutionary scenarios (section 3). Estimates of evolvability, $I_A$, and nucleotide diversity, $\pi$, were therefore log-transformed prior to analysis. Whilst true evolvabilities cannot be negative, measurement error can produce estimates below zero. To retain these data, we first tried left-censored models, treating non-positive values as censored between $-\infty$ and -15.903 on the log-scale, where -15.903 is the log of the minimum positive value reported. However, these models fitted the data poorly. We also tried a Gaussian model with log-link using the package 'rstan' v2.32.7 (20) to accommodate negative estimates on the data scale whilst allowing a multiplicative underlying model. However, issues with chain convergence prevented useful estimation of effects and variance components. Consequently, we finally chose to simply remove non-positive estimates for the final model. Given that only a small portion of evolvability estimates were non-positive (4.54%), the exclusion of these estimates should only have a minor quantitative effect on the results of the analyses and is not expected to impact our qualitative conclusions.

True variation in $ln(I_A)$ among species ($n = 108$), was estimated by fitting $ln(\hat{\pi})$ as a covariate with regression coefficient $\beta$, and *phylogenetic* and *non-phylogenetic* species effects as random. The phylogenetic covariance structure was assumed to be proportional to the amount of time two species have shared ancestry, based on a phylogeny constructed from published divergence-time estimates (Figure 2; 21). The tree was scaled to unit length such that the estimated variance in the phylogenetic ($V_{ln(I_A):\text{Phy}}$) and non-phylogenetic ($V_{ln(I_A):\text{S}}$) species effects sum to give the total between-species variation in $ln(I_A)$, after accounting for $ln(\hat{\pi})$. Differences in $ln(\hat{I_A})$ due to estimation methodology were accounted for with fixed effects for *method* and the number of *fixed* and *random effect terms*. We did not incorporate a formal measurement error structure as this would require the exclusion of all evolvability estimates lacking a reported or calculable measure of precision (25.89% of estimates for the largest $I_A$ model), limiting the precision with which model effects can be estimated. Trait differences in $ln(I_A)$ were modelled with fixed effects for *trait category* and *trait dimension* and a random effect for *trait* identity. A random *publication* effect was fitted to capture remaining differences among studies not explained by these predictors. True among-population variance was not estimated because few species had estimates from multiple populations across multiple publications, providing little power to partition variance at these levels.

The estimated parameters and variance components are reported in Tables 1 and 2. The slope of the regression was $\beta = 0.127$ [-0.202 – 0.410] (P=0.397), therefore doubling $\hat{\pi}$ would only increase mean evolvability by 9.2% [-13.1 – 32.8]. Despite substantial variation in $ln(I_A)$ between-species (standard deviation (SD) of 1.533 [0.649 – 2.426], only 0.7% [-3.0 – 9.5] of this could be predicted by $ln(\hat{\pi})$ (note we carry the sign in the $R^2$ value). The best estimates suggest that phylogenetic relatedness explained the largest proportion of among-species variation in $ln(I_A)$ (92.5% [52.1 – 100.0]) whereas residual non-phylogenetic species differences accounted for only a small proportion (5.3% [0.0 – 44.4]), however credible intervals were wide. $ln(I_A)$ did not significantly depend on the types of relatives ($\chi^2_6 = 6.934$, P=0.327), the number of fixed effects (0.011 [-0.103 – 0.128], P=0.869) or the number of random effects (0.324 [-0.017 – 0.659], P=0.060). There were suggestive (19) and significant differences in $ln(I_A)$ among different trait categories ($\chi^2_4 = 11.509$, P=0.021) and dimensions ($\chi^2_5 = 29.717$, P<0.001) respectively, and there was substantial among-trait variation that was not captured by these broad classifications (SD = 1.055 [0.881 – 1.221]). There was still considerable among-publication variance after accounting for differences in estimation and trait choice (SD = 1.175 [0.957 – 1.414]), but it is unclear to what extent this represents remaining methodological differences versus true differences between populations since among-population variance was not separately estimated. There was substantial residual variation (SD = 1.362 [1.296 – 1.440]) and the residuals were slightly more leptokurtic than normal.

We next repeated the analysis, substituting $ln(\hat{\pi})$ for $ln(\hat{\pi}_N/\hat{\pi}_S)$. The slope of the regression was $\beta$ = 0.086 [-0.599 – 0.753] (P=0.802), therefore doubling $\hat{\pi}_N/\hat{\pi}_S$ only corresponds to an increase in evolvability by 6.1% [-37.5 – 64.6] and 0.1% [-7.0 – 23.0] of true variation in $ln(I_A)$ between species ($n = 60$) could be predicted by $ln(\hat{\pi}_N/\hat{\pi}_S)$.

Microsatellite diversity has historically been used to infer the amount of neutral genetic variation within species (22). In the prior study (4), only a small number of species had both microsatellite $H_e$ and $I_A$ estimates, which prevented precise inference of the relationship between these measures. Here, we find no clear association between $ln(I_A)$ and $ln(\hat{H}_e)$ across the 66 species with estimates of both measures. The slope of the regression was $\beta$ = -0.535 [-1.308 – 0.344] (P=0.237), therefore doubling $\hat{H}_e$ may substantially reduce mean evolvability, $2^\beta$ = -31.0% [-66.2 – 15.8], but credible intervals are wide. The $R^2$ was effectively zero (-0.9% [-8.7 – 1.3]). A parallel analysis restricted to 41 species where both $H_e$ and $I_A$ were estimated in the same population produced similarly uncertain results. The slope of the regression was $\beta$ = 0.144 [-1.071 – 1.493] (P=0.835), meaning that evolvability is only expected to increase marginally if $\hat{H}_e$ doubled ($2^\beta$ = 10.5% [-64.8 – 144.0]) and the $R^2$ was again effectively zero (0.1% [-13.3 – 18.7]).

##### Molecular genetic diversity vs. Heritability

Given the weak correlation between $I_A$ and $h^2$ (Fig 2; 2), we performed additional analyses to quantify the relationship between the three measures of molecular genetic variation and $h^2$. Although quantitative genetic theory predicts a log-log linear relationship between $V_A$ and genetic diversity, it is not clear whether this expectation holds when $V_A$ is variance-standardised (i.e., when expressed as heritability, $h^2$). $h^2$ has a low dynamic range because it is typically bounded between 0-1, meaning that the results are unlikely to differ appreciably with- or without log-transformation of $h^2$. Therefore, we did not log-transform heritability and non-positive values were retained (7.4% of estimates). The priors, MCMC specifications and model structure are otherwise identical to the univariate models described above. As with the evolvability models, the relationships between $h^2$ and each of $ln(\pi)$, $ln(\pi_N/\pi_S)$ and $ln(H_e)$ were assessed individually. Across all heritability models, we only report the $R^2$ since a log-log regression would be required to quantify the expected proportional change in $h^2$ if genetic diversity were doubled ($2^\beta$).

The parameters and variance components estimated in the univariate analysis of $h^2$ and $ln(\hat{\pi})$ are reported in Tables 3 and 4. The regression slope was $\beta$ =-0.021 [-0.054 – 0.015] (P=0.243), and despite moderate between-species ($n = 130$) variation in $h^2$ (standard deviation (SD) of 0.134 [0.073 – 0.284]), very little -2.7%

[-23.0 – 2.6] of this could be predicted by $ln(\hat{\pi})$. There was low power to partition the total among-species variation in $h^2$ between phylogenetic (42.1% [0.0 – 97.8]) and residual non-phylogenetic species components (50.6% [0.0 – 93.5]).

Substituting $ln(\hat{\pi})$ for $ln(\hat{\pi}_N/\hat{\pi}_S)$ yielded similar conclusions, with $\beta$ =0.017 [-0.040 – 0.077] (P=0.590) and $R^2$ = 0.6% [-6.3 – 23.6] ($n$ = 48 species).

The relationship between $h^2$ and microsatellite $ln(\hat{H}_e)$ was significantly positive, with $\beta$ =0.088 [0.065 – 0.109](P<0.5×10$^{-3}$) and $R^2$ = 3.9% [0.3 – 8.8]. The parallel analysis restricted to species where both $H_e$ and $h^2$ were estimated in the same population produced very similar results, with $\beta$ =0.092 [0.070 – 0.116] (P<0.5×10$^{-3}$) and $R^2$ = 2.2% [0.2 – 5.9]. However, consistent with the findings of (4), this association was largely driven by a single study on *Arabidopsis thaliana* (23). When data from this study were excluded (140 $h^2$ estimates from 12 populations), the estimated relationship was weaker with wider credible intervals, both when heterozygosity estimates from different populations were included ($\beta$ =-0.033 [-0.130 – 0.062], P=0.512 and $R^2$ = -0.5% [-10.3 – 4.4]) and excluded ($\beta$ =-0.019 [-0.152 – 0.136], P=0.799 and $R^2$ = -0.1% [-8.0 – 6.1]).

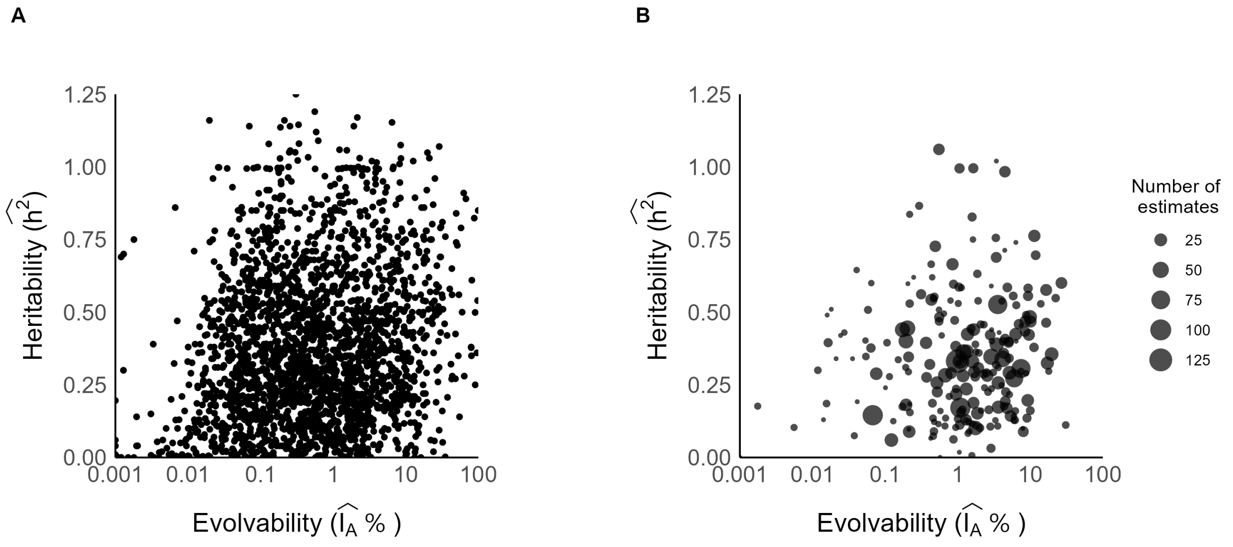


Figure 2: Measured heritability, $\widehat{h^2}$, against evolvability, $\widehat{I_A}$, for **(A)** 1,976 traits and **(B)** averaged across traits for 183 species, where the size of the point shows the number of estimates over which the average was calculated.

#### Bivariate models

In many species with estimates of quantitive genetic variation, estimates of molecular genetic diversity were not available. In order to leverage information from these species, we performed equivalent bivariate analyses for each of the molecular vs. quantitative genetic univariate models described above. In these models, both the trait-level quantitative genetic variation estimates ($ln(I_A)$ or $h^2$) and species-level molecular genetic diversity estimates ($ln(\pi)$, $ln(\pi_N/\pi_S)$ or $ln(H_e)$) were included as response variables. The model for the quantitative genetic response variable was equivalent to that of the respective univariate analysis but without molecular genetic diversity fitted as a predictor. For the molecular diversity response, variance among species was partitioned into a phylogenetic ($V_{M:Phy}$) and a non-phylogenetic 'residual' species ($V_{M:S}$) component. Correspondingly, the covariances between the two responses was estimated for the phylogenetic effects ($Cov_{Phy}$) and for the non-phylogenetic effects ($Cov_S$) and the overall species-level regression coefficient was obtained as

$$\beta = \frac{Cov_S + Cov_{Phy}}{V_{M:S} + V_{M:Phy}} \qquad (1)$$

Unlike the univariate analyses described in the main text, which implicitly assumes that the regression of quantitative genetic variation on molecular genetic diversity is the same at both phylogenetic and non-phylogenetic levels, the bivariate model allows the regression to be different. In the presence of phylogenetic effects, the bivariate model is expected to leverage additional information from species that only have estimates of quantitative genetic variation but for which estimates of molecular diversity exist for closely related taxa. While this may be expected to increase the precision with which the relationship between quantitative and molecular genetic variation can be estimated, this may be offset by allowing the regression at the two levels to differ rather than pooling information across these levels. Parameters relating solely to quantitative genetic variation are, however, expected to always be more precise. Priors and MCMC specifications are the same as for the univariate models except the prior for the species covariance matrix which was an inverse-Wishart distribution with the degree of belief parameter equal to 1.002 and a diagonal inverse scale matrix with 1.002 along the diagonal.

##### Molecular genetic diversity vs. Evolvability

The parameters estimated in the bivariate analysis of $ln(I_A)$ and $ln(\hat{\pi})$ (Tables 5 and 6) were broadly consistent with those of the univariate model. The association between $ln(V_A)$ and $ln(\hat{\pi})$ was again found to be weak ($\beta$ =-0.043 [-0.606 – 0.489], P=0.872; $2^\beta$ = -2.9% [-36.9 – 37.7] and $R^2$ =-0.2% [-20.3 – 15.8]). The between-species ($n$ = 193) variation in $ln(I_A)$ (standard deviation (SD) of 1.539 [0.874 – 2.401]) was mostly comprised of phylogenetic species effects (80.7% [52.4 – 95.9]), and residual non-phylogenetic species effects only accounted for a small proportion (12.3% [1.7 – 42.7]) of true variation among-species.

$ln(I_A)$ did not significantly differ among *types of relatives* ($\chi^2_6$ = 16.384, P=0.012), the number of *fixed effects* ( -0.081 [-0.169 – 0.021], P=0.087), or the number of *random effects* (0.102 [-0.106 – 0.356], P=0.369). $ln(I_A)$ significantly differed among different *trait categories* ($\chi^2_4$ = 25.287, P<0.001) and *dimensions* ($\chi^2_5$ = 49.659, P<0.001), and there was substantial among-trait variation that was not captured by these broad classifications (SD = 0.875 [0.725 – 1.016]). There was considerable among-publication variance after accounting for differences in estimation and trait choice (SD = 0.965 [0.772 – 1.136]) and substantial residual variation (SD = 1.409 [1.356 – 1.469]).

The association between $ln(I_A)$ and $ln(\hat{\pi}_N/\hat{\pi}_S)$ appeared stronger than in the univariate analysis, but remained too uncertain to be useful ($\beta$ =0.621 [-0.297 – 1.386], P=0.160; $2^\beta$ = 53.8% [-18.6 – 161.4] and $R^2$ =11.8% [-1.0 – 37.0]).

The relationship between $ln(I_A)$ and microsatellite $ln(\hat{H}_e)$ was again weak with wide credible intervals, both when diversity estimates from different populations were included ($\beta$ =-0.881 [-2.897 – 0.611], P=0.225; $2^\beta$ = -45.7% [-92.2 – 28.1] and $R^2$ = -4.3% [-27.5 – 3.4]), and excluded from the analysis ($\beta$ = -0.294 [-2.401 – 1.612], P=0.705; $2^\beta$ = -18.4% [-96.5 – 134.5] and $R^2$ = -0.3% [-25.1 – 7.3]).

##### Molecular genetic diversity vs. Heritability

The parameters and variance components estimated in the bivariate analysis of $h^2$ and $ln(\hat{\pi})$ are reported in Tables 7 and 8. The regression slope was $\beta$ =-0.009 [-0.038 – 0.015] (P=0.400), and little of the between-species ($n$ = 246) variation in $h^2$ (standard deviation (SD) of 0.162 [0.137 – 0.194]) could be predicted by $ln(\hat{\pi})$ ($R^2$ = -0.7% [-8.9 – 1.6]). Best estimates suggest that phylogenetic relatedness explained a small proportion of between-species variation in $h^2$ (3.0% [0.0 – 24.6]) and that residual between-species differences contributed substantially (95.2% [69.1 – 100.0]).

There was suggestive evidence that $h^2$ differs among types of relatives ($\chi^2_6$ = 15.41, P=0.017), and decreases with an increasing number of fixed (-0.013 [-0.023 – -0.003], P=0.008) and random effects (-0.026 [-0.048 – -0.003], P=0.031). $h^2$ significantly differed among different trait categories ($\chi^2_4$ = 23.503, P<0.001) but not dimensions ($\chi^2_5$ = 7.988, P=0.157), and there was considerable among-trait variation that was not captured by these broad classifications (SD = 0.078 [0.065 – 0.092]). There was substantial among-publication variance after accounting for differences in estimation and trait choice (SD = 0.134 [0.118 –

0.153]) and substantial residual variation (SD = 0.201 [0.196 – 0.207]).

For $ln(\hat{\pi}_N/\hat{\pi}_s)$, the regression slope was again weak ($\beta$ =-0.012 [-0.042 – 0.011], P=0.302) and variation in $ln(\hat{\pi}_N/\hat{\pi}_s)$ explained very little of the variation in $h^2$ ($R^2$ = -1.3% [-9.8 – 2.2]).

Likewise, models with microsatellite $ln(\hat{H}_e)$ included as the molecular diversity predictor showed no detectable association with $h^2$, both when diversity estimates obtained from different populations were included ($\beta$ =0.004 [-0.157 – 0.156], P=0.960 and $R^2$ = 0.0% [-6.4 – 7.0]) and excluded from the analyis ($\beta$ =0.014 [-0.128 – 0.157], P=0.812 and $R^2$ = 0.1% [-5.3 – 6.4]).

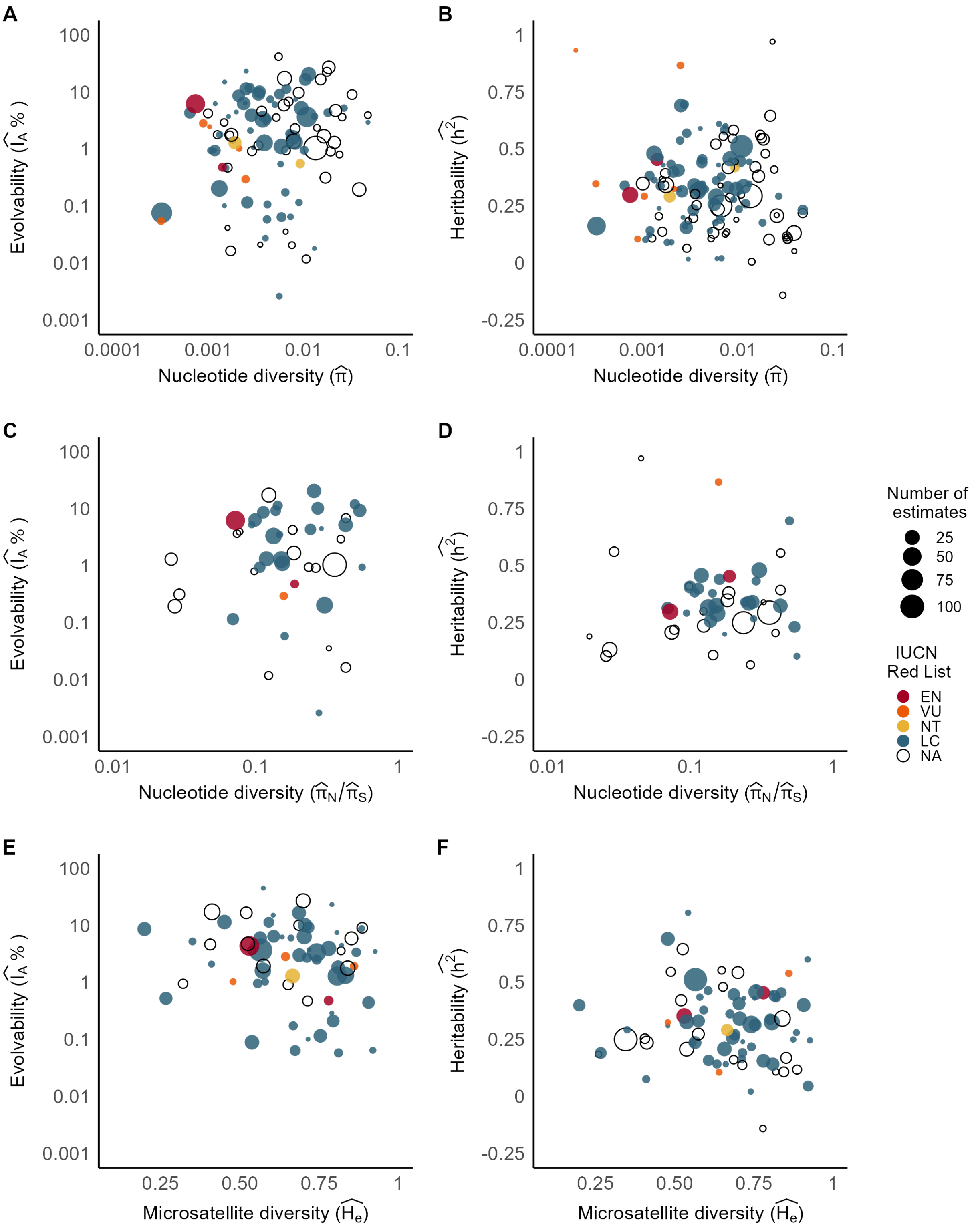


Figure 3: Measured species-mean quantitative genetic variation against molecular genetic variation: **(A)** Evolvability, $\widehat{I_A}$, and putatively neutral nucleotide diversity, $\hat{\pi}$ ($n$ =108); **(B)** Heritability, $\widehat{h^2}$, and putatively neutral nucleotide diversity, $\hat{\pi}$ ($n$ =130); **(C)** Evolvability and the ratio of nonsynonymous to synonymous nucleotide diversity, $\hat{\pi}_N/\hat{\pi}_S$ ($n$ =43); **(D)** Heritability and the ratio of nonsynonymous to synonymous nucleotide diversity ($n$ =48); **(E)** Evolvability and microsatellite expected heterozygosity, $\hat{H}_e$ ($n$ =66); **(F)** Heritability and microsatellite expected heterozygosity ($n$ =76).

### 3 Theoretical expectations

In this section, we outline the expected relationship between additive genetic variance ($V_A$) and neutral genetic diversity ($\pi$) under models of drift-selection-mutation balance. Throughout we assume additive effects on trait values and linkage-equilibrium. The models are covered in depth in Chapters 24 and 28 of 24.

#### Equilibrium: Under neutrality

Under mutation-drift balance the expected $V_A$ depends on the mutation model. Under the standard incremental model of mutation and additive effects (25),

$$V_A = 2N_eV_m \tag{2}$$

where $N_e$ is the effective population size and $V_m$ the mutational variance of the trait. Since the following models assume linkage-equilibrium it is typical to treat a locus as a region of length $L$ with mutation rate $L\mu$, where $\mu$ is the per-site mutation rate. Then, $V_m = 2nL\mu E[m^2]$ where $n$ is the number of regions contributing to the trait and $E[m^2]$ is the average squared effect of a new mutation on the trait. Under neutrality

$$\pi = 4N_e\mu/(1 + 4N_e\mu) \tag{3}$$

which is approximately $4N_e\mu$ when small, such that

$$\begin{aligned} ln(V_A) = & \quad ln(4N_e\mu) + ln(LnE[m^2]) \\ = & \quad ln(\pi) + ln(L) + ln(n) + ln(E[m^2]) \end{aligned} \tag{4}$$

and we expect a one-to-one relationship between $ln(V_A)$ and $ln(\pi)$ unless $\pi$ strongly covaries with $n$ and/or $E[m^2]$ over species/populations.

#### Equilibrium: Under stabilising selection

At the other extreme, we have models of mutation-selection balance in populations large enough that the effects of genetic drift are negligible. Here, the introduction of new mutational variance is counteracted by the strength of stabilising selection acting on genotypic values, denoted $1/V_S$. $V_S = \omega^2 + V_E$ where $\omega$

is the standard deviation of a Gaussian fitness function and $V_E$ is the environmental variation of the trait (typically set to one without loss of generality). Under continuum of alleles models there are two approximate solutions for the equilibrium additive genetic variance. When selection is strong relative to mutation we have the House-of-Cards approximation (26):

$$V_A = 4nL\mu V_S \tag{5}$$

a result that is in agreement with earlier biallelic models (27) with a domain of applicability of $20L\mu \leq E[m^2]/V_S$. Under pure stabilising selection, where the trait mean is at the optimum, this domain of applicability translates into $10L\mu \leq -E[s]$, where $E[s]$ is the average selection coefficient on a new mutation which is approximately $-E[m^2]/2V_S$ (28). Under this model,

$$ln(V_A) = ln(4L) + ln(\mu) + ln(n) + ln(V_S) \tag{6}$$

Here the regression of $ln(V_A)$ on $ln(\pi)$,

$$\beta = \frac{COV(ln(V_A), ln(\pi))}{VAR(ln(\pi))}, \tag{7}$$

reduces to

$$\beta = \frac{VAR(ln(\mu)) + COV(ln(\mu), ln(N_e))}{VAR(ln(\mu)) + VAR(ln(N_e)) + 2COV(ln(\mu), ln(N_e))} \tag{8}$$

again under the assumption that $n$ and $V_S$ are independent of $N_e$ and $\mu$. If all of the variation in $\pi$ is driven by differences in mutation rate, then $\beta = 1$ as in the neutral case. However, when there is substantial variation in $N_e$, $\beta < 1$ and populations/species with higher $\pi$ will not show proportionally higher $V_A$.

When mutation is strong relative to selection we have the Gaussian approximation (29):

$$\begin{aligned} V_A &= \sqrt{2nV_SV_m} \\ &= \sqrt{4n^2V_SL\mu E[m^2]} \end{aligned} \tag{9}$$

Under this model,

$$
\begin{aligned}
ln(V_A) = &\quad log(\sqrt{4n^2 V_S L \mu E[m^2]}) \\
= &\quad \tfrac{1}{2} log(4n^2 V_S L \mu E[m^2]) \\
= &\quad \tfrac{1}{2} log(4L) + log(n) + \tfrac{1}{2} ln(\mu) + \tfrac{1}{2} ln(V_S) + \tfrac{1}{2} ln(E[m^2])
\end{aligned} \tag{10}
$$

Following a similar logic as above

$$
\beta = \frac{1}{2} \frac{VAR(ln(\mu)) + COV(ln(\mu), ln(N_e))}{VAR(ln(\mu)) + VAR(ln(N_e)) + 2COV(ln(\mu), ln(N_e))} \tag{11}
$$

and $\beta = \frac{1}{2}$ when all of the variation in $\pi$ is driven by differences in mutation rate.

In conclusion, in extremely large populations, the relationship between $\pi$ and $V_A$ is expected to be weak if variation in mutation rate is small relative to variation in $N_e$. However, if variation in mutation rate is very large relative to variation in $N_e$ then the regression should lie between 1/2 (when selection is weak compared to mutation) and one (when selection is strong compared to mutation).

Models of mutation-selection-drift balance in populations that are small enough for genetic drift to have non-negligible effects produce outcomes intermediate between the neutral case and the pure mutation-selection models. However, deriving analytical results for the expected regression coefficient is more difficult because the equilibrium $V_A$ is not log-linear in $ln(N_e)$ (or $ln(\mu)$ under the Gaussian approximation). Instead, we can derive the partial derivatives of $ln(V_A)$ with respect to $ln(\mu)$ and $ln(N_e)$ to quantify how strongly $V_A$ responds to proportional changes in these individual components, and subsequently gain insight into the scenarios where $V_A$ is expected to scale strongly with $\pi$. When $ln(V_A)$ is linearly related to $ln(\mu)$ and $ln(N_e)$ and/or $ln(\mu)$ and $ln(N_e)$ are multivariate normal, the expected regression coefficients for $ln(\mu)$ and $ln(N_e)$ are equal to the expected partial derivatives over the distribution of $ln(\mu)$ and $ln(N_e)$ (30). If $ln(\mu)$ and $ln(N_e)$ do not vary too much, then the actual regression coefficients will not deviate too much from these expectations, or from the partial derivatives evaluated at the mean values of $ln(\mu)$ and $ln(N_e)$. Then, the expected regression coefficient is approximately:

$$
\begin{aligned}
\beta \approx &\quad \frac{COV(ln(\mu)\frac{\partial ln(V_A)}{\partial ln(\mu)}\Big|_{\overline{ln(\mu)}} + ln(N_e)\frac{\partial ln(V_A)}{\partial ln(N_e)}\Big|_{\overline{ln(N_e)}}, ln(\mu)+ln(N_e))}{VAR(ln(\mu))+VAR(ln(N_e))+2COV(ln(\mu),ln(N_e))} \\
&\quad \frac{VAR(ln(\mu))\frac{\partial ln(V_A)}{\partial ln(\mu)}\Big|_{\overline{ln(\mu)}} + VAR(ln(N_e))\frac{\partial ln(V_A)}{\partial ln(N_e)}\Big|_{\overline{ln(N_e)}} + COV(ln(N_e),ln(\mu))\left[\frac{\partial ln(V_A)}{\partial ln(\mu)}\Big|_{\overline{ln(\mu)}} + \frac{\partial ln(V_A)}{\partial ln(N_e)}\Big|_{\overline{ln(N_e)}}\right]}{VAR(ln(\pi))}
\end{aligned} \tag{12}
$$

where $\overline{ln(\mu)}$ and $\overline{ln(N_e)}$ are the average values of $ln(\mu)$ and $ln(N_e)$. When $ln(\mu)$ and $ln(N_e)$ are uncorrelated this reduces to

$$\beta \approx \frac{VAR(ln(\mu))\frac{\partial ln(V_A)}{\partial ln(\mu)}\Big|_{\overline{ln(\mu)}} + VAR(ln(N_e))\frac{\partial ln(V_A)}{\partial ln(N_e)}\Big|_{\overline{ln(N_e)}}}{VAR(ln(\pi))}. \tag{13}$$

Note that the correlation between $ln(\mu)$ and $ln(N_e)$ is most likely negative (31) in which case the actual regression will be shallower than this equation predicts since both partial derivatives are expected to be non-negative.

When selection is strong relative to mutation we have the House-of-Cards approximation (32):

$$V_A = \frac{4nL\mu V_S}{1+V_S/(N_eE[m^2])} \tag{14}$$

therefore, the partial derivative with respect to $ln(\mu)$ is simply one

$$\frac{\partial ln(V_A)}{\partial ln(\mu)} = 1 \tag{15}$$

For $ln(N_e)$ we have

$$\frac{\partial ln(V_A)}{\partial ln(N_e)} = \frac{\frac{V_S}{E[m^2]}}{N_e+\frac{V_S}{E[m^2]}} \tag{16}$$

which is a logistic function with a value of one as $N_e$ tends to zero, a growth rate of minus one and a midpoint (i.e. when $N_e = 1$) of $ln(V_S/E[m^2])$. Consequently, at low $N_e$ a one-to-one relationship is expected between $ln(V_A)$ and $ln(\pi)$ since both quantities scale with $\mu$ and $N_e$. The range of $N_e$ over which the relationship between $ln(V_A)$ and $ln(\pi)$ remains strong is dictated by $V_S/E[m^2]$. Specifically, when $4N_e < V_S/E[m^2]$ the partial derivative with respect to $ln(N_e)$ exceeds 0.8 (Figure 4). Note that this inequality corresponds to the scenario where $|4N_eE[s]| < 1$ and most new mutations are effectively neutral, as expected (24).

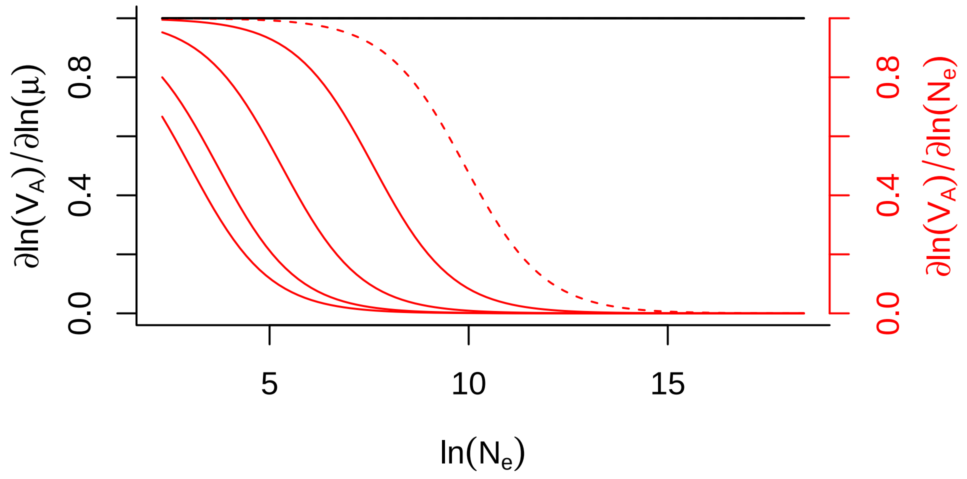


Figure 4: Partial derivatives of $ln(V_A)$ with respect to $ln(\mu)$ (black) and $ln(N_e)$ (red) as a function of $ln(N_e)$ under the House-of-Cards approximation where selection is strong relative to mutation. The derivative with respect to $ln(N_e)$ only depends on $V_S/E[m^2]$ which we set to 5, 10, 50, 500 and 5,000 corresponding to red lines from left to right. Under pure stabilising selection, these values correspond to average selection coefficients of -0.100, -0.050, -0.010, -0.001 and -0.0001 respectively. Solid lines indicate parameter combinations where the House-of-cards approximation is expected to hold ($20L\mu \leq E[m^2]/V_S$) and dashed lines not.

When mutation is relatively weak we have the Gaussian approximation (28):

$$V_A = \sqrt{\left(\frac{nV_S - 4n^2N_eL\mu E[m^2]}{2(N_e+n)}\right)^2 + \frac{4n^2N_eL\mu E[m^2]V_S}{N_e+n}} - \frac{nV_S - 4n^2N_eL\mu E[m^2]}{2(N_e+n)} \quad (17)$$

Note that we use the original derivation in (28) rather than the similar but simpler approximation in 24 (Equation 28.32c) which induces unintended behaviour in the derivatives. Nevertheless, the partial derivatives under the original derivation in 28 are ugly. With respect to $ln(\mu)$ the partial derivative is

$$\frac{\partial ln(V_A)}{\partial ln(\mu)} = 1 - \frac{V_m N_e \left(V_S - Q\right)}{SQ} \quad (18)$$

and for $ln(N_e)$ the derivative is

$$\frac{\partial ln(V_A)}{\partial ln(N_e)} = P\left(1 + \frac{N_e}{S}\left[V_m + \frac{V_S}{2n} - \frac{V_m V_S}{Q}\right]\right) \quad (19)$$

where

$$A = \frac{V_S}{2} - V_m N_e, \quad S = \sqrt{A^2 + 2V_m N_e V_S / P}, \quad Q = P(S + A) \text{ and } P = \frac{n}{N_e + n}. \tag{20}$$

In Figure 5 we plot these derivatives assuming $V_S = 20$ and $V_m = 0.001$ (33) but for a range of selection coefficients determined by $n$. When $N_e$ and/or selection coefficients are small the relationship between $ln(V_A)$ and $ln(\mu)$ is one, as seen under the House-of-Cards approximation. Only under strong selection or very large population sizes does the relationship between $ln(V_A)$ and $ln(\mu)$ tend to a half, as in the deterministic Gaussian approximation. However, such scenarios probably lie outside the domain of applicability of the Gaussian approximation (since mutation is no longer strong relative to selection), so in practice the relationship between $ln(V_A)$ and $ln(\mu)$ is likely to equal one. As with the House-of-Cards approximation, the relationship between $ln(V_A)$ and $ln(N_e)$ is one when $N_e$ is small and tends to zero as $N_e$ and/or selection coefficients increase in magnitude. However, with the Gaussian approximation, $V_A$ remains dependent on $N_e$ over a greater range of values; for instance, the $V_A$ of a highly polygenic trait under weak stabilising selection may show strong dependence on $N_e$ if $N_e < 20,000$.

In conclusion, a strong one-to-one relationship between $ln(V_A)$ and $ln(\pi)$ is only expected when quantitative traits are neutral. If they are under stabilising selection, any relationship between $ln(V_A)$ and $ln(\pi)$ is likely driven by their shared dependence on the mutation rate and so the relationship is probably weak if the main contribution to variation in $\pi$ is variation in $N_e$. However, if $N_e$ is very small and/or the strength of selection on mutations that affect quantitative traits is very weak, then the relationship between $ln(V_A)$ and $ln(\pi)$ also has a contribution from their shared dependence on $N_e$ and the relationship may be stronger. However, the dependence of $V_A$ on $N_e$ is quickly lost if $N_e$ exceeds a few hundred, at least under the House-of-cards approximation where selection is assumed strong relative to mutation. While the Gaussian approximation predicts that this dependence remains strong for $N_e$ of a few thousand, the House-of-cards approximation has historically been viewed to be more accurate for most systems (26). However, it is not clear whether this view remains valid in light of recent data (e.g. 35).

#### Non-equilibrium: Under neutrality

The above results apply to populations at equilibrium. In non-equilibrium populations, such as those that have undergone a recent bottleneck or expansion, it has been suggested that the dynamics of $\pi$ and $V_A$ are

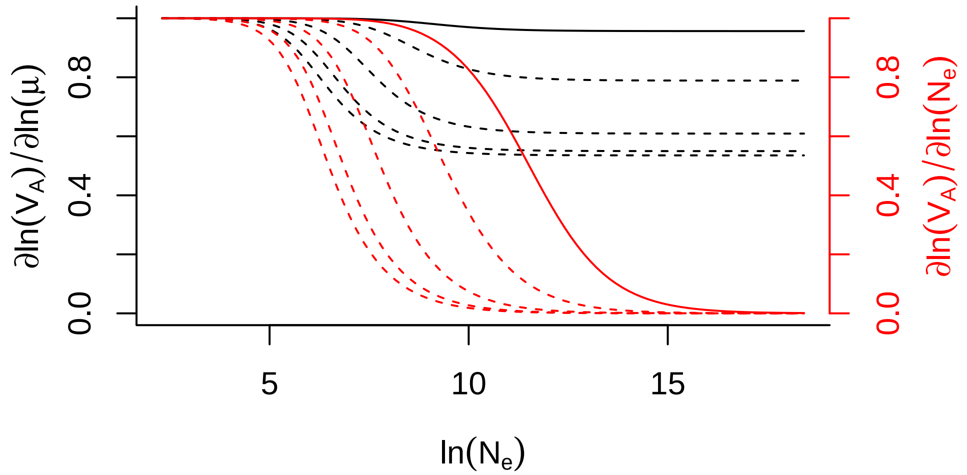


Figure 5: Partial derivatives of $ln(V_A)$ with respect to $ln(\mu)$ (black) and $ln(N_e)$ (red) as a function of $ln(N_e)$ under the Gaussian approximation where selection is weak relative to mutation. The derivatives depend in complicated ways on $V_S$, $V_m$ and $n$. In all cases $V_S = 20$ and $V_m = 0.001$. However, we vary $n$ from 100, 200, 1000, 10000 and 100000 corresponding to red and black lines from left to right. Given a mutation rate of $10^8$ and $L$=1,000 these values correspond to selection coefficients of -0.100, -0.050, -0.010, -0.001 and -0.0001, respectively. Solid lines indicate parameter combinations where the deterministic (i.e $N_e = \infty$) Gaussian approximation is expected to hold ($20L\mu \geq E[m^2]/V_S$) and dashed lines not. However, the domain of applicability of the Gaussian approximation is expected to be smaller when drift is higher (34).

on sufficiently different time scales that the two measures may be weakly related, even in small populations (36, 37). To explore these non-equilibrium dynamics, we use recurrence equations which describe how the value of $V_A$ or $\pi$ at generation $t$ depends on its value in the previous generation, $t-1$. The recurrence equation for $V_A$ assuming the trait is neutral is (25)

$$V_A(t) = \quad V_A(t-1)\left(1 - \frac{1}{2N_e(t-1)}\right) + V_m \tag{21}$$

For a more extensive treatment with dominance see (38). The recurrence equation for $\pi$ is (39):

$$\begin{aligned} \pi(t) = &\quad 1 - (1-\mu)^2\left[1 - \pi(t-1)\left(1 - \frac{1}{2N_e(t-1)}\right)\right] \\ \pi(t) = &\quad (1-\mu)^2\pi(t-1)\left(1 - \frac{1}{2N_e(t-1)}\right) + \mu(2-\mu) \end{aligned} \tag{22}$$

When $\mu$ and $\pi$ are small, terms in $\mu^2$ and $\mu\pi$ can be ignored, giving

$$\pi(t) = \pi(t-1)\left(1 - \frac{1}{2N_e(t-1)}\right) + 2\mu \tag{23}$$

If at time $t-1$, $V_A(t-1) = \pi(t-1)V_m/2\mu$, then

$$\begin{aligned} V_A(t) &= \frac{V_m}{2\mu}\pi(t-1)\left(1 - \frac{1}{2N_e(t-1)}\right) + V_m \\ V_A(t) &= \frac{V_m}{2\mu}\left(\pi(t-1)\left(1 - \frac{1}{2N_e(t-1)}\right) + 2\mu\right) \\ V_A(t) &= \frac{V_m}{2\mu}\pi(t) \end{aligned} \tag{24}$$

such that $V_A(t) \propto \pi(t)$ in generations thereafter (assuming $V_m$ and $\mu$ remain constant) and so the relationship between $ln(V_A(t))$ and $ln(\pi(t))$ will be one-to-one. While the condition $V_A(t-1) = \pi(t-1)V_m/2\mu$ seems restrictive, note that it is satisfied at equilibrium under neutrality since $\pi = 4N_e\mu$ and $V_A = 2V_mN_e$ such that $V_A = \pi V_m/2\mu$. Consequently, under neutrality a population only has to come to equilibrium once for the relationship between $ln(V_A(t))$ and $ln(\pi(t))$ to remain close to one-to-one. This relationship persists even if the population later departs from equilibrium due to changes in population size, and is only disrupted by external perturbations to allele frequencies (e.g. a translocation event). To illustrate this idea, in Figure 6 we iterate Equations 21 and 22 for a thousand generations with $N_e = 100$ and $\pi(0) = 0.01$. We do this for four parameter combinations with $V_m$ equal to 0.001 or 0.0001 and $\mu$ equal to $10^{-8}$ or $10^{-6}$. Since $\pi(0) = 0.01$ is greater than what would be expected given $N_e = 100$ and the mutation rates, $\pi(0)$ is not currently at equilibrium and so $\pi$ is expected to decrease over time. For each parameter combination, two populations were iterated. One where $V_A$ was initiated at its expected value given $\pi(0)$ ($V_A(0) = \pi(0)V_m/2\mu$: dashed lines) and one where $V_A$ initiated at one (solid lines). In all populations, $V_A(0)$ is higher than what would be expected given values for $V_m$ and $N_e = 100$. However, simulations where $V_A(0)$ is initiated at $\pi(0)V_m/2\mu$ are consistent with a scenario where $V_A$ and $\pi$ had come to their equilibrium at some previous point in time where $N_e$ was higher. We refer to these populations as previously equilibrated. While we see non-linearity between $ln(V_A)$ and $ln(\pi)$ when sampled (over time) from populations that were never previously equilibrated, populations that were previously equilibrated show close to linear one-to-one relationships.

#### Non-equilibrium: Under stabilising selection

Selected loci are expected to equilibrate faster than neutral loci: in the extreme case lethals equilibrate in one generation. Consequently, when a trait is under selection, $V_A$ may equilibrate at a faster rate than $\pi$. The recurrence equation under the Gaussian approximation is (40):

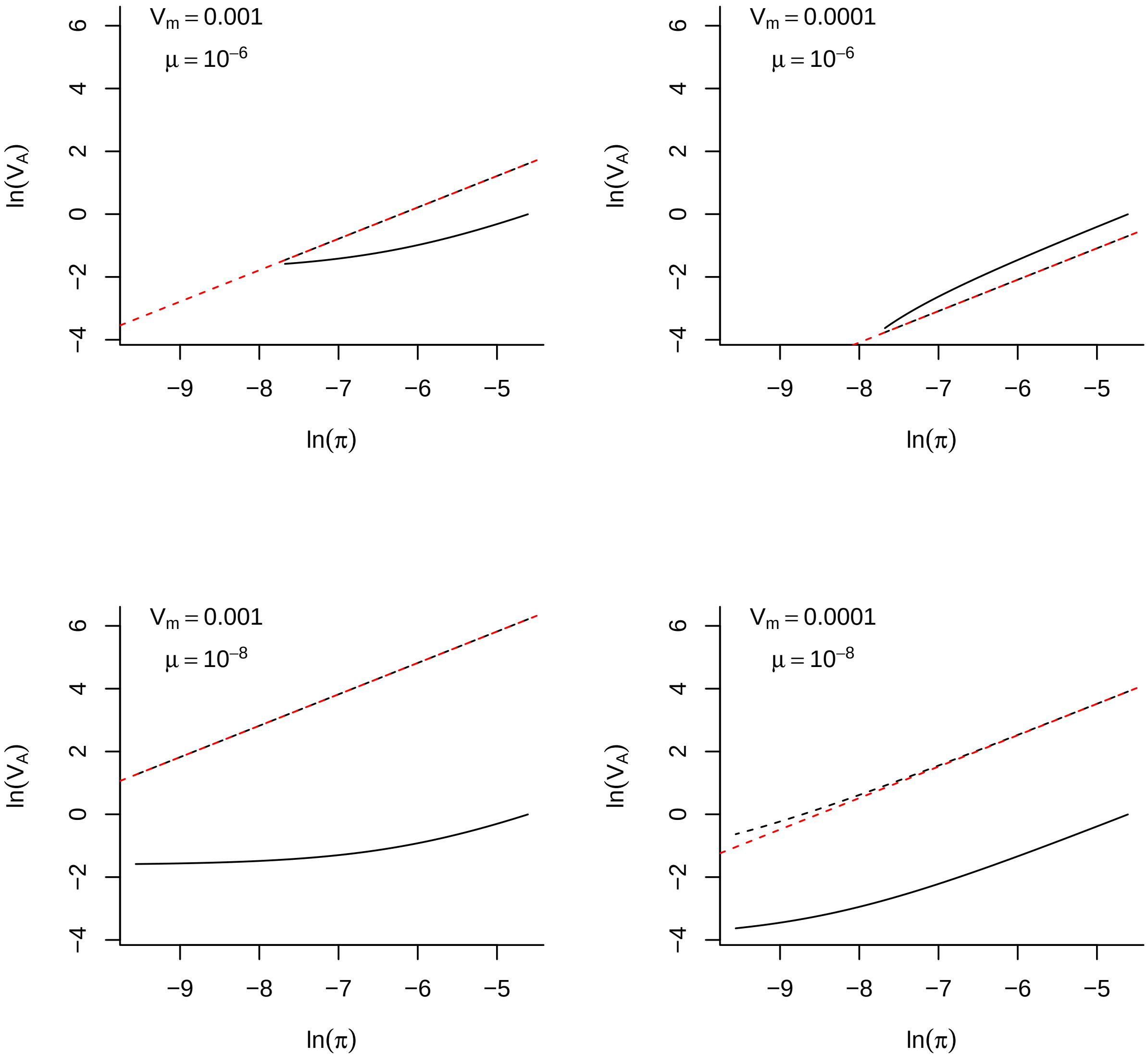


Figure 6: Populations are initialised with a genetic diversity, $\pi(0)$, of 0.01. Solid black lines are for populations where the additive genetic variance, $V_A(0)$, was initialised at one (not previously equilibrated) and the dashed black lines are for previously equilibrated populations where $V_A(0) = \pi(0)V_m/2\mu$. Note that in all cases all quantities are far above their expected equilibrium values given $N_e = 100$ and so the quantities are moving from right to left over time, representing the loss of genetic variation following a bottleneck. Nevertheless, in previously equilibrated populations, $log(V_A)$ has a close to linear dependence on $log(\pi)$ with a slope of one and intercept of $log(V_m/2\mu)$ as would be seen in permanently equilibrated populations (red dashed lines).

$$V_A(t) = V_A(t-1)\left[1 - \frac{1}{2N_e(t-1)} + \frac{1}{2n}\left(1 - \frac{1}{N_e(t-1)}\right)\frac{V_A(t-1)}{V_A(t-1)+V_S}\right] + V_m$$

(25)

and as $V_S$ increases (i.e. the strength of stabilising selection diminishes) or $n$ increases (i.e. the strength of stabilising selection per region diminishes) the recurrence equation tends to the neutral case. Note that the original derivation (40) is in terms of $kh^2$ where $k$ is the proportional reduction in the phenotypic variance caused by stabilising selection and $h^2$ is the trait heritability: $kh^2 = V_A/(V_A + V_S)$. Setting $V_A(t-1) = \pi(t-1)V_m/2\mu$ as before, we have

$$\begin{aligned} V_A(t) = & \ \tfrac{V_m}{2\mu}\left(\pi(t-1)\left[1 - \tfrac{1}{2N_e(t-1)}\right] + 2\mu + \pi(t-1)\tfrac{1}{2n}\left(1 - \tfrac{1}{N_e(t-1)}\right)\tfrac{V_A(t-1)}{V_A(t-1)+V_S}\right) \\ V_A(t) = & \ \tfrac{V_m}{2\mu}\left(\pi(t) + \pi(t-1)\tfrac{1}{2n}\left(1 - \tfrac{1}{N_e(t-1)}\right)\tfrac{V_A(t-1)}{V_A(t-1)+V_S}\right) \end{aligned} \quad (26)$$

which shows that $V_A$ will equilibrate faster than $\pi$ when the strength of stabilising selection is strong and/or there are few loci contributing to $V_A$. In Figure 7 we initiate populations at their equilibrium values for $\pi$ (Equation 3) and $V_A$ (Equation 17) given an effective population size of 1,000 and then follow their trajectory following a bottleneck event where $N_e$ instantaneously becomes 100. As in the neutral scenario above, we use four parameter combinations with $V_m$ equal to 0.001 or 0.0001 and $\mu$ equal to $10^{-8}$ or $10^{-6}$. For each parameter combination we iterate Equations 26 and 22 when the number of loci is either 100 (selection per locus is weak: solid black lines) or 10 (selection per locus is strong: solid red lines). When selection per locus is weak the relationship is close to linear with a slope of one reflecting the neutral case, but when selection per locus is strong the relationship between $ln(V_A)$ and $ln(\pi)$ becomes weaker, particularly if $V_m$ is large.

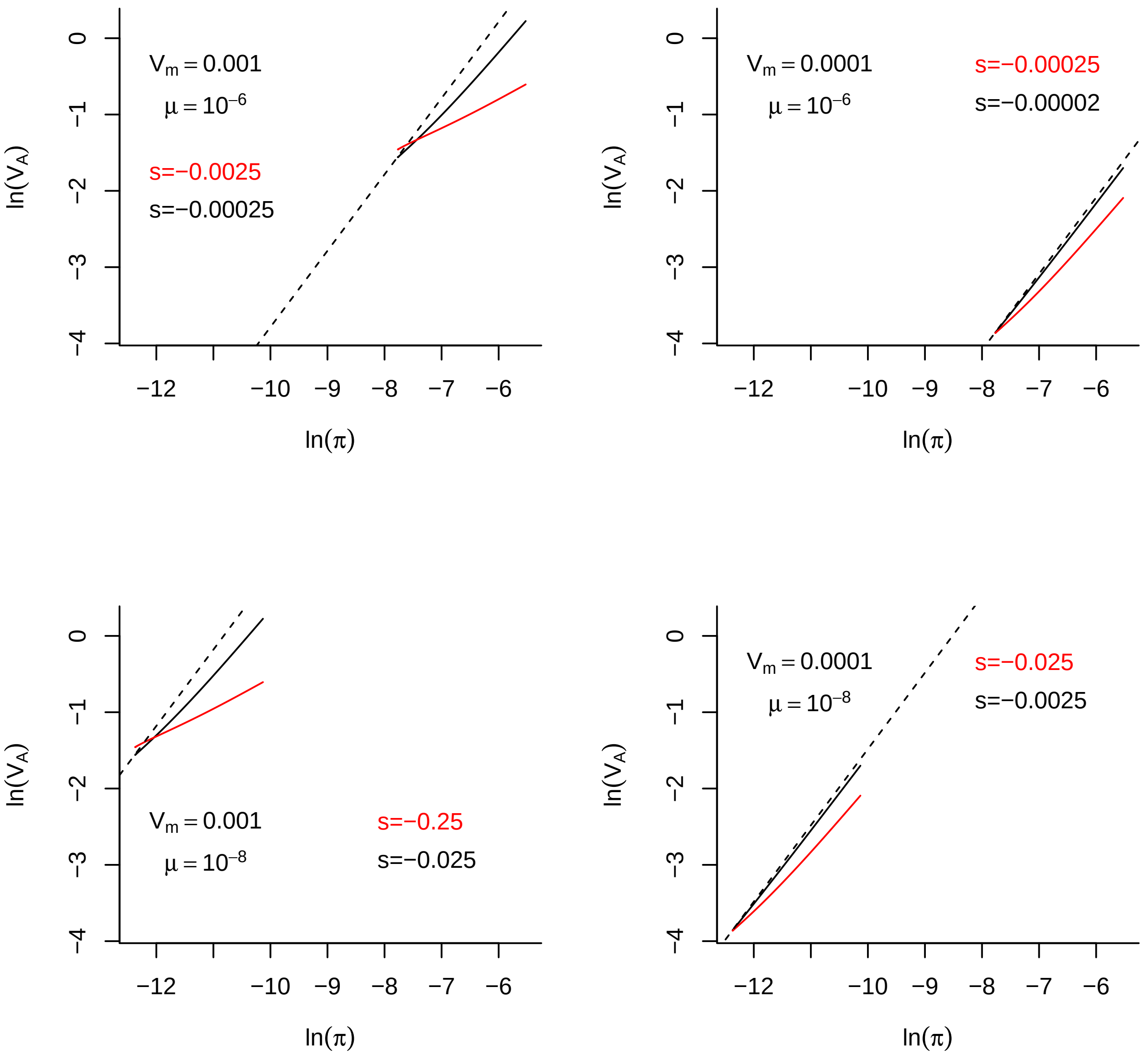


Figure 7: Populations are initialised with a genetic diversity ($\pi(0)$) equal to the neutral expectation when $N_e = 1,000$. Similarly, the additive genetic variance is initialised ($V_A(0)$) at its Gaussian mutation-selection-drift balance expectation. Following initialisation, populations experience a bottleneck event where $N_e$ instantaneously becomes 100. We follow their trajectory when the number of loci contributing to the trait is large ($n = 100$) and so selection per-locus is weak (solid black line) and when the number of loci is small ($n = 10$) and so selection per-locus is strong (solid red line). Different scenarios involve different combinations of $V_m$ and $\mu$ but $V_s = 20$ and $L$=1,000 are constant. Note that in the neutral case these initial conditions would give rise to a relationship that is close to one-to-one with an intercept of $ln(V_m/2\mu)$ (dashed black line - See Figure 6).

### 4 Supplementary Tables

Table 1: Summary of fixed effects (posterior mean, lower and upper 95% credible intervals) from the univariate analysis of evolvability, $ln(I_A)$. pMCMC is twice the posterior probability that the effect is less than or greater than zero (whichever is smaller) and $P(>\chi^2)$ is the P-value of an omnibus test (a Wald test using the posterior mean and covariance matrix) for multi-category factors (displayed next to the final level). The intercept corresponds to linear morphological traits measured using a pedigree-based (animal model) estimate of relatedness.

| Parameter | mean | l-95% | u-95% | pMCMC | $P(>\chi^2)$ |
|---|---|---|---|---|---|
| Intercept | -0.303 | -2.829 | 2.251 | 0.789 | |
| ln($\pi$) | 0.130 | -0.202 | 0.410 | 0.397 | |
| Trait type: fitness | -0.712 | -2.232 | 0.722 | 0.369 | |
| Trait type: life history | 0.123 | -0.596 | 0.824 | 0.748 | |
| Trait type: behaviour | 1.472 | 0.525 | 2.398 | 0.003 | |
| Trait type: physiological | 0.320 | -0.243 | 0.807 | 0.225 | 0.021 |
| Method: mid-parent-offspring | -1.040 | -2.085 | -0.127 | 0.031 | |
| Method: full-sib | 0.110 | -0.853 | 1.040 | 0.809 | |
| Method: half-sib | 0.233 | -0.457 | 1.045 | 0.537 | |
| Method: clonal | 0.319 | -0.720 | 1.269 | 0.545 | |
| Method: realized | -0.431 | -2.260 | 1.252 | 0.625 | |
| Method: single-parent-offspring | 0.068 | -0.764 | 0.903 | 0.871 | 0.327 |
| n.fixed | 0.009 | -0.103 | 0.128 | 0.869 | |
| n.random | 0.323 | -0.017 | 0.659 | 0.060 | |
| Dimension: quadratic | 0.736 | -0.034 | 1.453 | 0.054 | |
| Dimension: cubic | 0.988 | 0.461 | 1.471 | $<0.5\times10^{-3}$ | |
| Dimension: meristic | 1.361 | 0.730 | 2.018 | $<0.5\times10^{-3}$ | |
| Dimension: time | 0.322 | -0.451 | 1.156 | 0.441 | |
| Dimension: other | 0.454 | -0.129 | 0.972 | 0.107 | $<0.001$ |

Table 2: Summary of estimated variance components (posterior mode, lower and upper 95% credible intervals) from the univariate analysis of evolvability, $ln(I_A)$, across 108 species.

| | **Standard deviation** | | | **% of between-species variance** | | | **% of total variance** | | |
|---|---|---|---|---|---|---|---|---|---|
| **Variance component** | mode | l-95% | u-95% | mode | l-95% | u-95% | mode | l-95% | u-95% |
| ln($\pi$) | 0.051 | $0.116\times10^{-3}$ | 0.404 | 0.060 | $0.433\times10^{-6}$ | 8.303 | 0.005 | $0.192\times10^{-6}$ | 2.325 |
| Phylogenetic | 1.229 | 0.434 | 2.414 | 99.388 | 52.055 | 99.981 | 23.814 | 6.268 | 59.253 |
| Non-phylogenetic species | 0.400 | $0.845\times10^{-3}$ | 0.781 | 0.237 | $0.106\times10^{-4}$ | 44.444 | 0.078 | $0.342\times10^{-5}$ | 10.134 |
| Trait | 1.015 | 0.881 | 1.221 | | | | 15.626 | 9.210 | 24.381 |
| Publication | 1.205 | 0.957 | 1.414 | | | | 20.458 | 10.921 | 32.572 |
| Residual | 1.355 | 1.296 | 1.440 | | | | 26.437 | 14.933 | 37.151 |
| Between-species | 1.571 | 0.649 | 2.426 | | | | 29.352 | 13.373 | 61.997 |
| Total | 2.470 | 2.172 | 3.223 | | | | | | |

Table 3: Summary of fixed effects (posterior mean, lower and upper 95% credible intervals) from the univariate analysis of heritability, $h^2$. pMCMC is twice the posterior probability that the effect is less than or greater than zero (whichever is smaller) and $P(>\chi^2)$ is the P-value of an omnibus test (a Wald test using the posterior mean and covariance matrix) for multi-category factors (displayed next to the final level). The intercept corresponds to linear morphological traits measured using a pedigree-based (animal model) estimate of relatedness.

| Parameter | mean | l-95% | u-95% | pMCMC | $P(>\chi^2)$ |
|---|---|---|---|---|---|
| Intercept | 0.275 | 0.014 | 0.534 | 0.020 | |
| ln($\pi$) | -0.021 | -0.054 | 0.015 | 0.243 | |
| Trait type: fitness | -0.145 | -0.315 | 0.034 | 0.110 | |
| Trait type: life history | -0.042 | -0.097 | 0.012 | 0.132 | |
| Trait type: behaviour | -0.105 | -0.182 | -0.024 | 0.010 | |
| Trait type: physiological | -0.055 | -0.100 | -0.004 | 0.024 | 0.019 |
| Method: mid-parent-offspring | -0.010 | -0.112 | 0.090 | 0.863 | |
| Method: full-sib | 0.095 | $0.562\times10^{-3}$ | 0.190 | 0.051 | |
| Method: half-sib | 0.021 | -0.062 | 0.107 | 0.651 | |
| Method: clonal | 0.079 | -0.012 | 0.184 | 0.117 | |
| Method: realized | 0.150 | -0.006 | 0.319 | 0.066 | |
| Method: single-parent-offspring | 0.074 | -0.011 | 0.150 | 0.073 | 0.193 |
| n.fixed | -0.008 | -0.019 | 0.005 | 0.234 | |
| n.random | -0.034 | -0.064 | -0.006 | 0.017 | |
| Dimension: quadratic | 0.034 | -0.048 | 0.123 | 0.455 | |
| Dimension: cubic | 0.011 | -0.045 | 0.063 | 0.693 | |
| Dimension: meristic | 0.007 | -0.052 | 0.065 | 0.799 | |
| Dimension: time | 0.010 | -0.056 | 0.084 | 0.806 | |
| Dimension: other | -0.027 | -0.069 | 0.012 | 0.193 | 0.412 |

Table 4: Summary of estimated variance components (posterior mode, lower and upper 95% credible intervals) from the univariate analysis of heritability, $h^2$. across 130 species

| | Standard deviation | | | % of between-species variance | | | % of total variance | | |
|---|---|---|---|---|---|---|---|---|---|
| **Variance component** | mode | l-95% | u-95% | mode | l-95% | u-95% | mode | l-95% | u-95% |
| ln($\pi$) | 0.030 | $0.299\times10^{-4}$ | 0.056 | 0.145 | $0.7\times10^{-5}$ | 21.868 | 0.024 | $0.981\times10^{-6}$ | 3.576 |
| Phylogenetic | 0.007 | $0.121\times10^{-3}$ | 0.275 | 0.622 | $0.933\times10^{-4}$ | 97.839 | 0.223 | $0.172\times10^{-4}$ | 50.493 |
| Non-phylogenetic species | 0.090 | 0.024 | 0.131 | 0.803 | $0.245\times10^{-3}$ | 93.461 | 10.171 | $0.152\times10^{-3}$ | 17.237 |
| Trait | 0.095 | 0.080 | 0.112 | | | | 9.441 | 5.186 | 14.757 |
| Publication | 0.141 | 0.120 | 0.168 | | | | 22.538 | 11.421 | 30.648 |
| Residual | 0.201 | 0.195 | 0.208 | | | | 48.212 | 26.787 | 55.453 |
| Between-species | 0.125 | 0.073 | 0.284 | | | | 16.629 | 7.160 | 53.399 |
| Total | 0.296 | 0.270 | 0.388 | | | | | | |

Table 5: Summary of estimated fixed effects (posterior mean, lower and upper 95% credible intervals) from the bivariate analysis of evolvability, $ln(I_A)$. Nucleotide diversity, $ln(\pi)$, was included as a response but had no fixed predictors. pMCMC is twice the posterior probability that the effect is less than or greater than zero (whichever is smaller) and $P(>\chi^2)$ is the P-value of an omnibus test (a Wald test using the posterior mean and covariance matrix) for multi-category factors (displayed next to the final level). The intercept for $log(I_A)$ corresponds to linear morphological traits measured using a pedigree-based (animal model) estimate of relatedness.

| Parameter | mean | l-95% | u-95% | pMCMC | $P(>\chi^2)$ |
|---|---|---|---|---|---|
| Intercept: ln($\pi$) | -0.873 | -2.555 | 0.913 | 0.291 | |
| Intercept: ln($I_A$) | -5.166 | -6.377 | -3.818 | $<0.5\times10^{-3}$ | |
| Trait type: fitness | -0.379 | -1.837 | 1.097 | 0.620 | |
| Trait type: life history | 0.099 | -0.500 | 0.681 | 0.730 | |
| Trait type: behaviour | 1.686 | 1.023 | 2.441 | $<0.5\times10^{-3}$ | |
| Trait type: physiological | 0.178 | -0.300 | 0.623 | 0.465 | $< 0.001$ |
| Method: mid-parent-offspring | -1.276 | -2.030 | -0.442 | $<0.5\times10^{-3}$ | |
| Method: full-sib | 0.475 | -0.216 | 1.184 | 0.192 | |
| Method: half-sib | 0.222 | -0.334 | 0.774 | 0.425 | |
| Method: clonal | 0.277 | -0.521 | 1.086 | 0.497 | |
| Method: realized | -0.649 | -2.011 | 0.612 | 0.347 | |
| Method: single-parent-offspring | -0.185 | -0.806 | 0.525 | 0.598 | 0.012 |
| n.fixed | -0.082 | -0.169 | 0.021 | 0.087 | |
| n.random | 0.104 | -0.106 | 0.356 | 0.369 | |
| Dimension: quadratic | 0.691 | 0.018 | 1.382 | 0.059 | |
| Dimension: cubic | 1.086 | 0.663 | 1.500 | $<0.5\times10^{-3}$ | |
| Dimension: meristic | 1.355 | 0.829 | 1.857 | $<0.5\times10^{-3}$ | |
| Dimension: time | 0.265 | -0.387 | 0.863 | 0.421 | |
| Dimension: other | 0.685 | 0.211 | 1.155 | 0.005 | $< 0.001$ |

Table 6: Summary of estimated variance components (posterior mode, lower and upper 95% credible intervals) from the bivariate analysis of evolvability, $ln(I_A)$, across 193 species. Nucleotide diversity, $ln(\pi)$, was included as a response, and the proportion of variation in $ln(I_A)$ attributable to $ln(\pi)$ was calculated from their (co)variance.

| | **Standard deviation** | | | **% of between-species variance** | | | **% of total variance** | | |
|---|---|---|---|---|---|---|---|---|---|
| **Variance component** | mode | l-95% | u-95% | mode | l-95% | u-95% | mode | l-95% | u-95% |
| ln($\pi$) | 0.110 | $0.111\times10^{-3}$ | 0.873 | 0.102 | $0.507\times10^{-6}$ | 19.226 | 0.036 | $0.244\times10^{-6}$ | 9.464 |
| Phylogenetic | 1.094 | 0.595 | 2.235 | 88.236 | 52.447 | 95.926 | 21.693 | 10.071 | 56.286 |
| Non-phylogenetic species | 0.498 | 0.332 | 0.796 | 5.966 | 1.696 | 42.698 | 3.251 | 1.091 | 10.962 |
| Trait | 0.843 | 0.725 | 1.016 | | | | 12.288 | 6.677 | 18.428 |
| Publication | 0.959 | 0.772 | 1.136 | | | | 14.880 | 7.308 | 23.713 |
| Residual | 1.407 | 1.356 | 1.469 | | | | 35.534 | 20.947 | 44.818 |
| Between-species | 1.226 | 0.874 | 2.401 | | | | 44.427 | 18.281 | 62.776 |
| Total | 2.365 | 2.086 | 3.039 | | | | | | |

Table 7: Summary of estimated fixed effects (posterior mean, lower and upper 95% credible intervals) from the bivariate analysis of heritability, $h^2$. Nucleotide diversity, $ln(\pi)$, was included as a response but had no fixed predictors. pMCMC is twice the posterior probability that the effect is less than or greater than zero (whichever is smaller) and $P(>\chi^2)$ is the P-value of an omnibus test (a Wald test using the posterior mean and covariance matrix) for multi-category factors (displayed next to the final level). The intercept for $h^2$ corresponds to linear morphological traits measured using a pedigree-based (animal model) estimate of relatedness.

| Parameter | mean | l-95% | u-95% | pMCMC | $P(>\chi^2)$ |
|---|---|---|---|---|---|
| Intercept: ln($\pi$) | 0.376 | 0.304 | 0.449 | $<0.5\times10^{-3}$ | |
| Intercept: $h^2$) | -4.860 | -6.140 | -3.528 | $<0.5\times10^{-3}$ | |
| Trait type: fitness | -0.164 | -0.310 | -0.016 | 0.031 | |
| Trait type: life history | -0.033 | -0.074 | 0.010 | 0.124 | |
| Trait type: behaviour | -0.092 | -0.155 | -0.033 | 0.004 | |
| Trait type: physiological | -0.080 | -0.120 | -0.043 | $<0.5\times10^{-3}$ | $< 0.001$ |
| Method: mid-parent-offspring | 0.034 | -0.050 | 0.108 | 0.415 | |
| Method: full-sib | 0.114 | 0.038 | 0.192 | 0.003 | |
| Method: half-sib | $-0.466\times10^{-3}$ | -0.068 | 0.061 | 0.976 | |
| Method: clonal | 0.079 | -0.008 | 0.161 | 0.062 | |
| Method: realized | 0.146 | 0.026 | 0.279 | 0.030 | |
| Method: single-parent-offspring | 0.071 | 0.010 | 0.135 | 0.034 | 0.017 |
| n.fixed | -0.013 | -0.023 | -0.003 | 0.008 | |
| n.random | -0.026 | -0.048 | -0.003 | 0.031 | |
| Dimension: quadratic | 0.002 | -0.066 | 0.082 | 0.959 | |
| Dimension: cubic | -0.011 | -0.054 | 0.030 | 0.642 | |
| Dimension: meristic | -0.022 | -0.065 | 0.022 | 0.322 | |
| Dimension: time | $0.13\times10^{-4}$ | -0.054 | 0.049 | 0.978 | |
| Dimension: other | -0.038 | -0.070 | -0.008 | 0.018 | 0.157 |

Table 8: Summary of estimated variance components (posterior mode, lower and upper 95% credible intervals) from the bivariate analysis of heritability, $h^2$, across 246 species. Nucleotide diversity, $ln(\pi)$, was included as a response, and the proportion of variation in $h^2$ attributable to $ln(\pi)$ was calculated from their (co)variance.

| | **Standard deviation** | | | **% of between-species variance** | | | **% of total variance** | | |
|---|---|---|---|---|---|---|---|---|---|
| **Variance component** | mode | l-95% | u-95% | mode | l-95% | u-95% | mode | l-95% | u-95% |
| ln($\pi$) | 0.002 | $0.544\times10^{-4}$ | 0.050 | 0.040 | $0.76\times10^{-5}$ | 7.761 | 0.016 | $0.294\times10^{-5}$ | 2.600 |
| Phylogenetic | 0.006 | $0.117\times10^{-4}$ | 0.091 | 0.158 | $0.567\times10^{-6}$ | 24.630 | 0.027 | $0.155\times10^{-6}$ | 8.441 |
| Non-phylogenetic species | 0.152 | 0.135 | 0.177 | 99.384 | 69.114 | 99.997 | 25.717 | 20.707 | 32.669 |
| Trait | 0.079 | 0.065 | 0.092 | | | | 6.770 | 4.602 | 9.173 |
| Publication | 0.130 | 0.118 | 0.153 | | | | 19.470 | 15.229 | 25.003 |
| Residual | 0.200 | 0.196 | 0.207 | | | | 43.628 | 38.693 | 49.193 |
| Between-species | 0.161 | 0.137 | 0.194 | | | | 29.265 | 22.029 | 36.970 |
| Total | 0.297 | 0.286 | 0.320 | | | | | | |

Table 9: Summary of fixed effects (posterior mean, lower and upper 95% credible intervals) from the univariate analysis of evolvability, $ln(I_A)$, with IUCN Red List status included as a predictor. pMCMC is twice the posterior probability that the effect is less than or greater than zero (whichever is smaller) and $P(>\chi^2)$ is the P-value of an omnibus test (a Wald test using the posterior mean and covariance matrix) for multi-category factors (displayed next to the final level). The intercept corresponds to linear morphological traits measured using a pedigree-based (animal model) estimate of relatedness.

| Parameter | mean | l-95% | u-95% | pMCMC | $P(>\chi^2)$ |
|---|---|---|---|---|---|
| Intercept | -1.226 | -3.597 | 1.274 | 0.245 | |
| IUCN: near threatened | 0.223 | -0.993 | 1.264 | 0.668 | |
| IUCN: vulnerable | -0.103 | -1.134 | 1.066 | 0.865 | |
| IUCN: endangered | -0.023 | -1.305 | 1.206 | 0.975 | |
| IUCN: critically endangered | 0.310 | -1.039 | 1.702 | 0.668 | 0.992 |
| Trait type: fitness | 0.207 | -1.363 | 1.911 | 0.827 | |
| Trait type: life history | 0.198 | -0.523 | 0.996 | 0.648 | |
| Trait type: behaviour | 1.694 | 0.931 | 2.532 | $<0.5\times10^{-3}$ | |
| Trait type: physiological | 0.250 | -0.304 | 0.854 | 0.372 | 0.002 |
| Method: mid-parent-offspring | -1.254 | -2.220 | -0.246 | 0.023 | |
| Method: full-sib | 0.826 | -0.255 | 1.903 | 0.148 | |
| Method: half-sib | 1.031 | -0.008 | 2.070 | 0.055 | |
| Method: clonal | 1.093 | -0.121 | 2.459 | 0.103 | |
| Method: realized | 2.448 | -0.178 | 4.937 | 0.062 | |
| Method: single-parent-offspring | 0.389 | -0.465 | 1.291 | 0.400 | 0.004 |
| n.fixed | 0.027 | -0.073 | 0.136 | 0.620 | |
| n.random | 0.247 | -0.013 | 0.490 | 0.047 | |
| Dimension: quadratic | 0.906 | 0.052 | 1.762 | 0.040 | |
| Dimension: cubic | 1.046 | 0.493 | 1.559 | $<0.5\times10^{-3}$ | |
| Dimension: meristic | 1.247 | 0.573 | 1.877 | $<0.5\times10^{-3}$ | |
| Dimension: time | -0.086 | -0.973 | 0.698 | 0.839 | |
| Dimension: other | 0.505 | -0.053 | 1.102 | 0.088 | $< 0.001$ |

Table 10: Summary of fixed effects (posterior mean, lower and upper 95% credible intervals) from the univariate analysis of heritability, $h^2$, with IUCN Red List status included as a predictor. pMCMC is twice the posterior probability that the effect is less than or greater than zero (whichever is smaller) and $P(>\chi^2)$ is the P-value of an omnibus test (a Wald test using the posterior mean and covariance matrix) for multi-category factors (displayed next to the final level). The intercept corresponds to linear morphological traits measured using a pedigree-based (animal model) estimate of relatedness.

| Parameter | mean | l-95% | u-95% | pMCMC | $P(>\chi^2)$ |
|---|---|---|---|---|---|
| Intercept | 0.319 | -0.056 | 0.603 | 0.077 | |
| IUCN: near threatened | -0.002 | -0.133 | 0.142 | 0.990 | |
| IUCN: vulnerable | 0.157 | 0.017 | 0.290 | 0.020 | |
| IUCN: endangered | 0.070 | -0.101 | 0.237 | 0.409 | |
| IUCN: critically endangered | 0.065 | -0.097 | 0.230 | 0.447 | 0.179 |
| Trait type: fitness | -0.137 | -0.288 | 0.045 | 0.119 | |
| Trait type: life history | -0.022 | -0.087 | 0.030 | 0.475 | |
| Trait type: behaviour | -0.076 | -0.149 | -0.006 | 0.041 | |
| Trait type: physiological | -0.107 | -0.152 | -0.063 | $<0.5\times10^{-3}$ | $< 0.001$ |
| Method: mid-parent-offspring | 0.003 | -0.089 | 0.091 | 0.956 | |
| Method: full-sib | 0.223 | 0.117 | 0.332 | $<0.5\times10^{-3}$ | |
| Method: half-sib | -0.001 | -0.122 | 0.122 | 0.991 | |
| Method: clonal | 0.245 | 0.122 | 0.387 | $<0.5\times10^{-3}$ | |
| Method: realized | 0.068 | -0.169 | 0.278 | 0.552 | |
| Method: single-parent-offspring | 0.095 | 0.022 | 0.167 | 0.005 | 0.001 |
| n.fixed | -0.009 | -0.020 | 0.002 | 0.096 | |
| n.random | -0.029 | -0.052 | -0.006 | 0.010 | |
| Dimension: quadratic | 0.006 | -0.081 | 0.114 | 0.921 | |
| Dimension: cubic | 0.002 | -0.049 | 0.053 | 0.939 | |
| Dimension: meristic | -0.034 | -0.091 | 0.025 | 0.242 | |
| Dimension: time | -0.005 | -0.073 | 0.064 | 0.872 | |
| Dimension: other | -0.026 | -0.067 | 0.009 | 0.178 | 0.671 |

Table 11: Number of species with estimates of evolvability, $I_A$, or heritability, $h^2$, grouped by IUCN Red List status

| | **Number of Species** | |
|---|---|---|
| **IUCN status** | **Evolvability** | **Heritability** |
| Least concern | 100 | 124 |
| Near threatened | 5 | 5 |
| Vulnerable | 7 | 10 |
| Endangered | 3 | 3 |
| Critically Endangered | 2 | 4 |
| Not classified | 76 | 100 |
